# Paralogous lncRNAs CYTOR and MORRBID share a conserved trans acting function in MEK ERK signaling

**DOI:** 10.64898/2026.06.24.734157

**Authors:** Tahleel Ali-Nasser, Susie Altalef Mishaan, Ornit Bhonkar, Zhongyang Lin, Youfen Qian, Nisrine Lahoud-Jeries, Dvir Aran, Assaf C. Bester

## Abstract

**Background:** Long non-coding RNAs (lncRNAs) exhibit rapid evolutionary turnover, often driven by genomic duplication. How paralogous lncRNAs maintain, partition, or diverge in function across distinct genomic contexts remains poorly understood. The evolutionarily conserved lncRNA MORRBID and its primate-specific paralog CYTOR provide a natural framework to interrogate the functional consequences of lncRNA duplication.

**Results:** Although CYTOR and MORRBID have acquired distinct transcript variants influenced by their divergent genomic environments, we demonstrate that they maintain a robust, shared core function encoded by near-identical dominant two-exon transcripts. Using SNP-based paralog-specific quantification, we found that CYTOR contributes more strongly to the shared transcript pool, while both transcripts localize predominantly to the cytoplasm, consistent with a shared trans-acting function. Simultaneous repression of CYTOR and MORRBID consistently impairs cell adhesion and migration across multiple cancer models. Mechanistically, the shared CYTOR/MORRBID transcript pool associates with MEK2 and sustains MEK-ERK signaling. This signaling axis promotes FOSL1 expression and AP-1-linked transcriptional output, including expression of the downstream effector EPHA4, whose role was supported by rescue experiments. Patient tumor transcriptomes and healthy single-cell datasets further supported the associated mesenchymal, adhesion, and epithelial-mesenchymal transition program.

**Conclusions:** Our findings establish that paralogous lncRNAs can retain a conserved mechanistic core despite context-dependent transcriptional divergence. The CYTOR/MORRBID transcript pool defines a shared lncRNA signaling module that supports MAPK-ERK signaling and adhesion-migration programs across cancer and mesenchymal-like cellular contexts. This defines a unified mechanistic framework for the shared core function of these widely studied paralogous lncRNAs.

## Background

Long non-coding RNAs (lncRNAs) are transcripts longer than 200 nucleotides that lack protein-coding potential and constitute the largest class of non-coding RNAs in humans^1^. Although numerous lncRNAs have been implicated in diverse biological processes, they are typically expressed at low levels and exhibit highly cell type-specific functions that are difficult to generalize across systems^2^. Moreover, the limited sequence conservation observed for most lncRNAs raises the possibility that many arise from pervasive transcription at regulatory DNA elements rather than representing bona fide functional molecules ^3–5^. These features; restricted expression, low abundance and poor conservation, highlight a central dilemma in the field which it, how to reconcile the expanding repertoire of reported lncRNA functions with properties that, by conventional criteria, argue against widespread biological significance. Resolving this discrepancy remains a major challenge in defining lncRNA function^1^.

This challenge is further complicated by the rapid and dynamic evolution of lncRNA loci. Compared with protein-coding genes (PCGs), lncRNAs frequently emerge de novo, diverge rapidly, or are lost during evolution^3,6–8^. While some lncRNAs exhibit conservation at the level of primary sequence, others are conserved only in structural features, genomic position (synteny), expression patterns, or functional output^5,6,9^. These observations suggest that lncRNA function may depend less on primary sequence than on genomic context and regulatory interactions^3,10,11^. However, how these evolutionary features contribute to the preservation, loss, or reconfiguration of lncRNA function remains unclear^3^. Paralogous lncRNAs provide a useful framework to address this question, as they enable shared ancestry to be examined alongside divergence in genomic context and regulatory environment.

Here, we address this question using the evolutionarily conserved lncRNA MIR4435-2HG (MORRBID) and its primate-specific paralog LINC00152 (CYTOR). MORRBID was first characterized in mouse as a regulator of the neighboring gene Bcl2l11 (Bim) through a transcript-dependent mechanism involving the recruitment of epigenetic silencers^12–15^. In contrast, CYTOR has been studied primarily in human systems, where it has been linked to cytoplasmic regulatory functions, most commonly through proposed miRNA-sponging mechanisms^16^. More broadly, the literature on MORRBID and CYTOR describes multiple context-dependent activities but lacks a unifying framework connecting their shared ancestry to underlying mechanisms^12–15,17–19^. Together, these observations suggest functional divergence despite a common evolutionary origin.

A segmental duplication event in primates gave rise to CYTOR, largely preserving promoter and gene sequence features while relocating the locus to a distinct genomic environment^13–15,20^. This evolutionary history provides a natural system to test how genomic context shapes lncRNA function ^3,10^. Although MORRBID and CYTOR retain related transcriptional features, they reside in distinct regulatory landscapes, raising the question of whether they have functionally diverged or whether shared biological activities can be maintained despite genomic relocation and lineage-specific evolution.

Here, we use MORRBID and CYTOR as a model to investigate how lncRNA paralogs acquire and maintain function across evolutionary divergence. Despite their distinct genomic contexts, we show that these lncRNAs retain shared functional outputs. Both regulate MAPK/ERK signalling through direct interaction with MEK2 in the cytoplasm, thereby modulating cell adhesion and migration in normal and cancer cells. Together, these findings demonstrate that lncRNA function can be preserved independently of genomic context and provide a framework for understanding how regulatory interactions shape lncRNA functional evolution.

## Results

### lncRNAs MORRBID and CYTOR are functionally indistinguishable

The long non-coding RNA (lncRNA) MORRBID (MIR4435-2HG/LINC00978) is located on human chromosome 2q13 and is positionally conserved in mice between BCL2L11 and ANAPC1^12^ (Fig.1a). In both species, MORRBID is transcribed from the negative strand. In primates, a ∼200-kilobase segmental duplication encompassing the MORRBID locus gave rise to the human lncRNA CYTOR (LINC00152) ^14^. In humans, this duplication relocated CYTOR to chromosome 2p11, where it is transcribed from the plus strand in a head-to-head configuration with the protein-coding gene RGPD2 (Fig.1a). Given their implication in health and disease ^18,21^, these paralogs provide a useful model for investigating the evolutionary origins and functional acquisition of lncRNAs. Both MORRBID and CYTOR are multi-exonic lncRNAs that give rise to several transcript variants (Fig. 1a). To assess how this segmental duplication influenced isoform expression, we analyzed RNA sequencing (RNA-seq) data from the Genotype-Tissue Expression (GTEx) project across 54 non-diseased tissues. For both loci, we identified a single dominant two-exon isoform (CYTOR: ENST00000331944; MORRBID: ENST00000409569) (Fig. 1b).

**Fig. 1.**
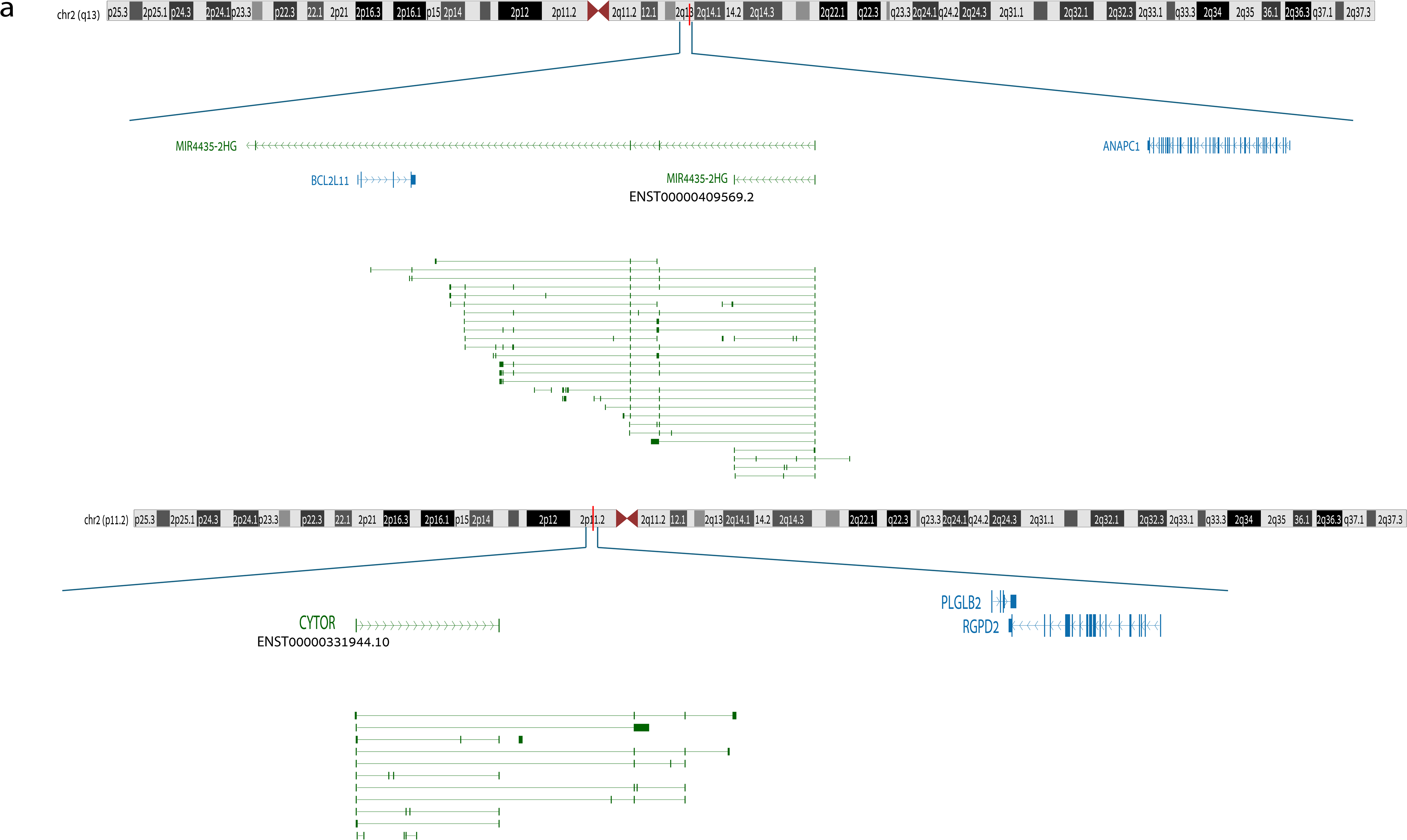
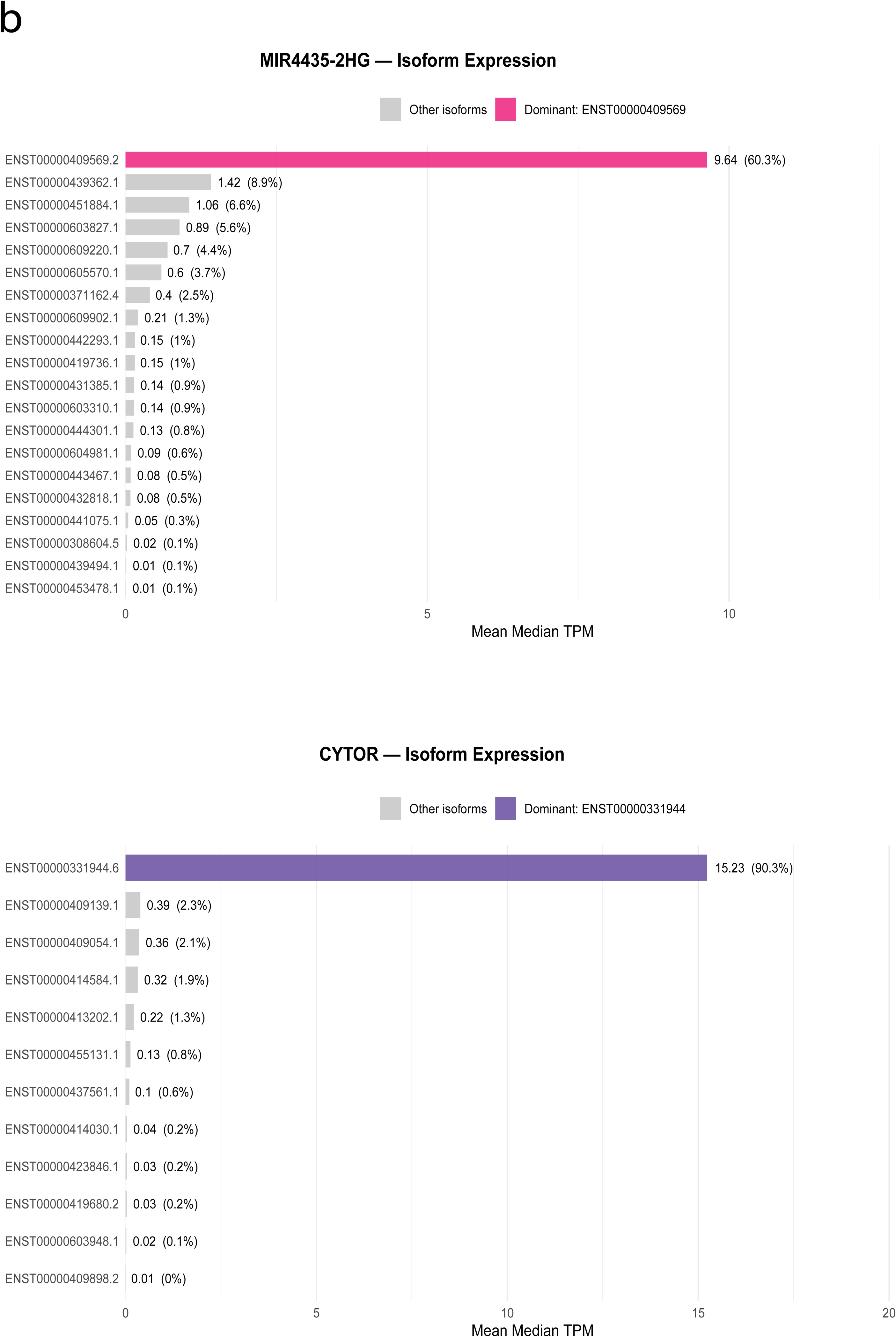
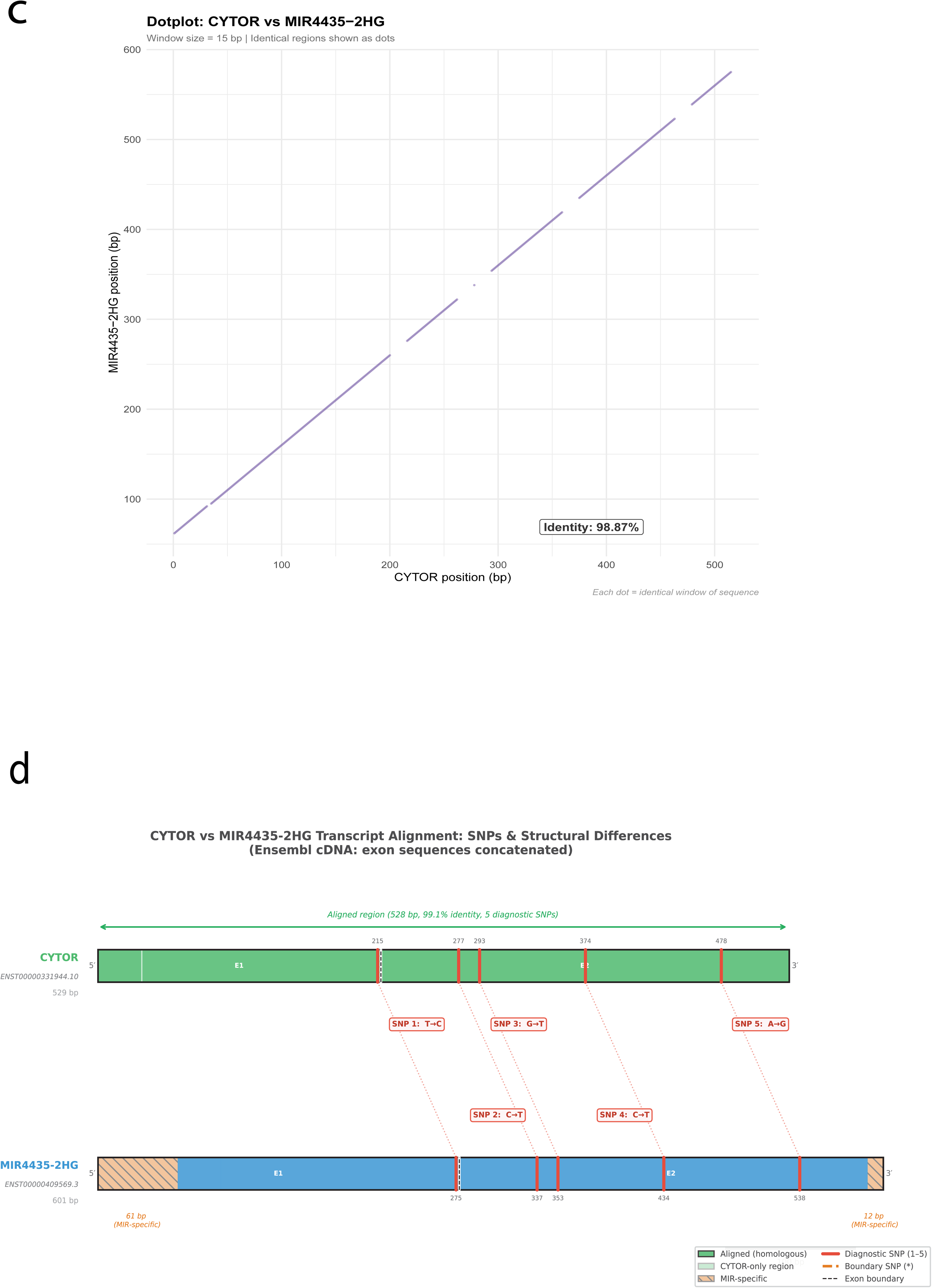
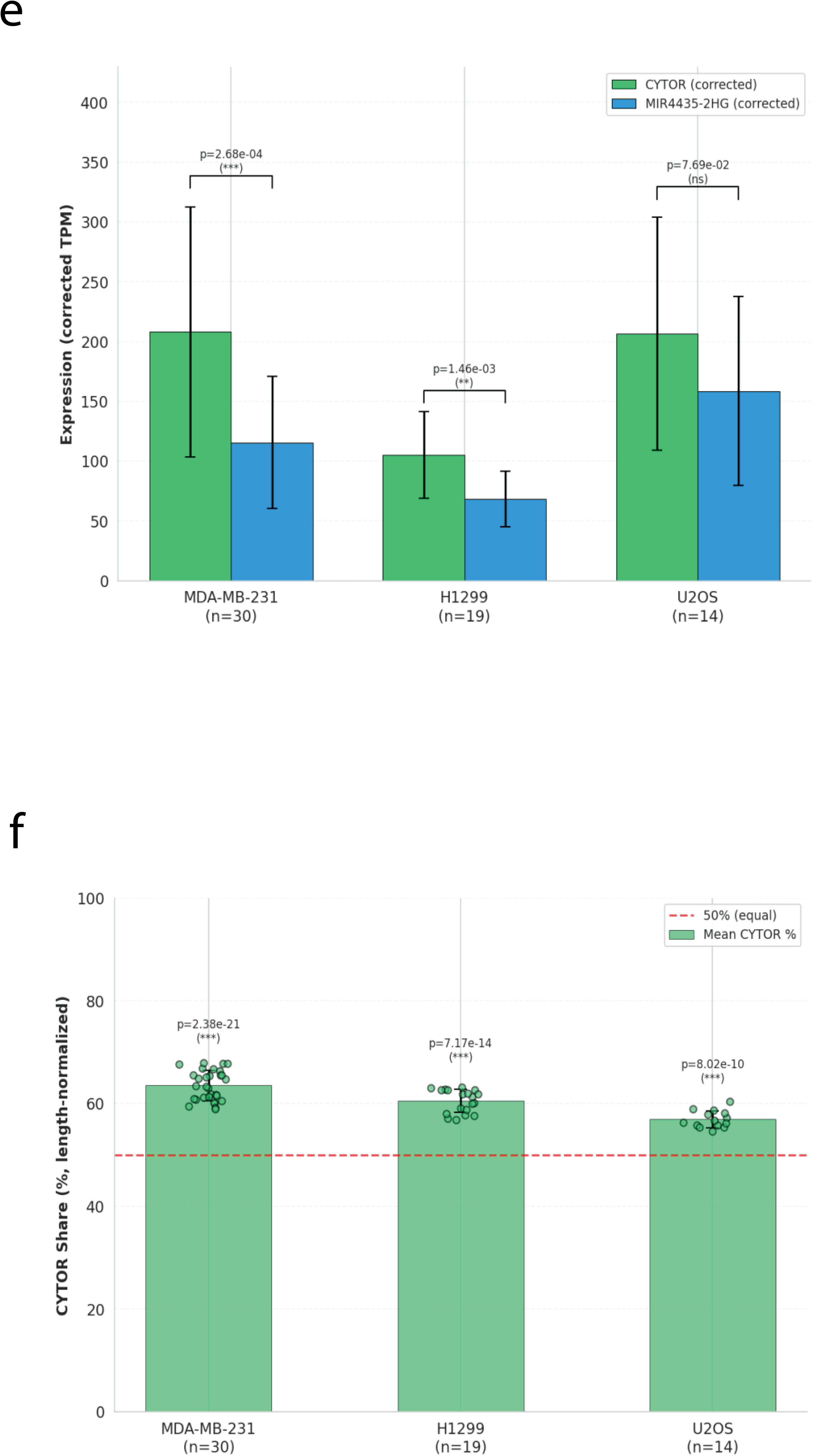
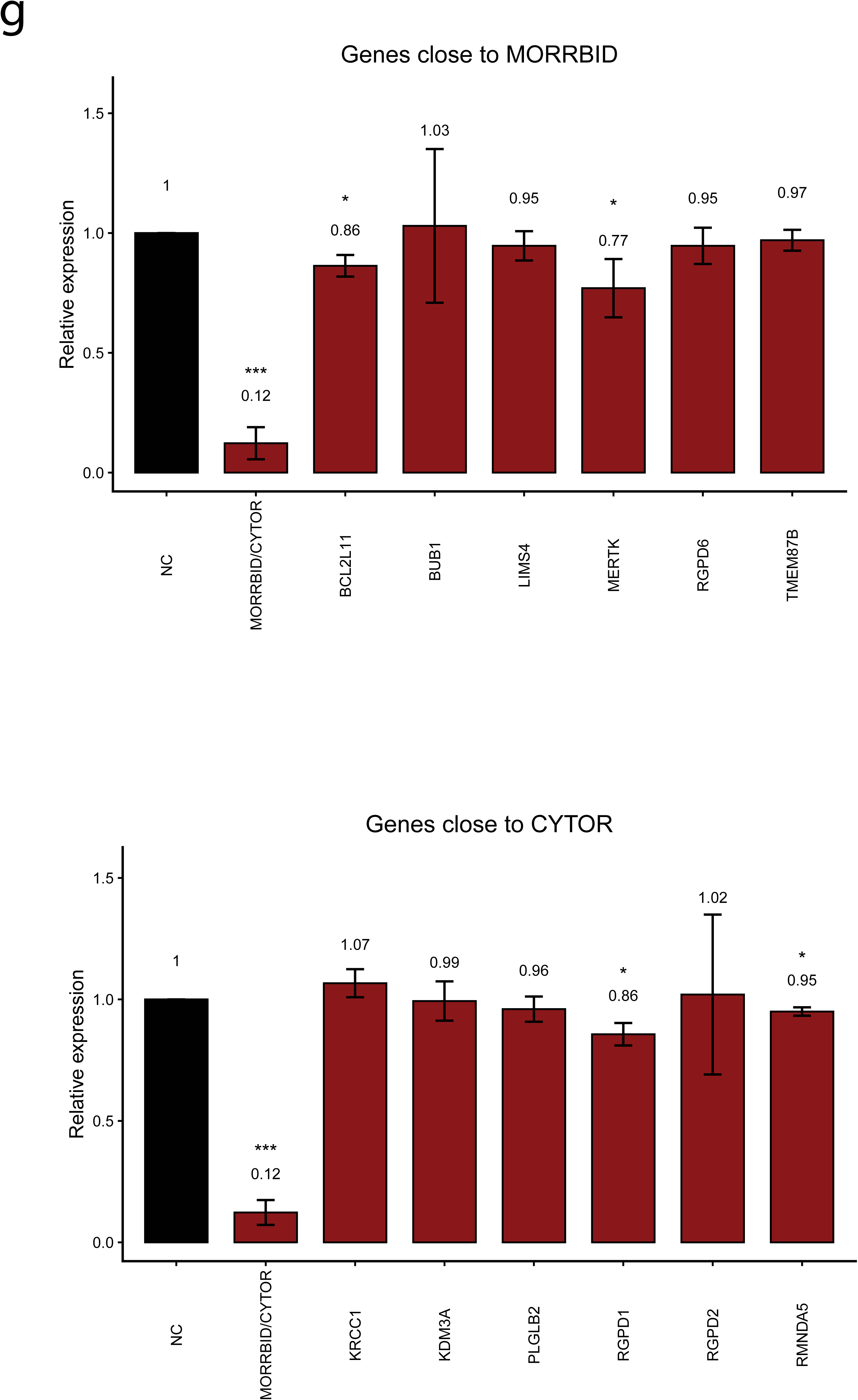
lncRNAs MORRBID and CYTOR are highly conserved paralogs with similar transcript architecture and expression profiles. **a** Schematic representation of the genomic loci of MORRBID and CYTOR on chromosome 2, showing their transcriptional orientation and neighboring protein-coding genes (PCGs). The positions of annotated transcript isoforms and adjacent genes are shown in the context of the surrounding chromosomal regions. Adapted from the UCSC Genome Browser. **b** Isoform expression profiles of MORRBID and CYTOR across GTEx v7 tissues showing dominant expression of a single two-exon isoform for each locus (MORRBID: ENST00000409569; CYTOR: ENST00000331944). Total transcript per million (TPM) values indicate mean median transcript abundance across tissues. **c** Dot-plot alignment of dominant CYTOR and MORRBID transcript isoforms (MORRBID: ENST00000409569; CYTOR: ENST00000331944) demonstrating extensive sequence conservation. **d** Pairwise alignment of dominant CYTOR and MORRBID transcript isoforms showing >98% nucleotide identity and five diagnostic single-nucleotide differences between transcripts. **e** SNP-corrected transcript abundance of CYTOR and MIR4435-2HG (MORRBID) in MDA-MB-231, H1299, and U2OS cell lines. TPM estimated by Salmon for the combined paralogous locus was redistributed according to the SNP-derived expression ratio to obtain paralog-specific corrected TPM values. Bars represent mean ± s.d.; P values were calculated using two-sided Mann-Whitney U tests. Sample sizes (n) are indicated below each group. **f** Length-normalized CYTOR transcript fraction derived from diagnostic SNP analysis in the indicated cell lines. The proportion of SNP-informative reads carrying CYTOR-specific alleles was quantified using a read-level multi-SNP classification approach and normalized for transcript length to estimate the relative contribution of CYTOR to total CYTOR + MIR4435-2HG expression. The dashed red line indicates equal expression (50%). Bars represent mean ± SD.; dots indicate individual biological replicates. P values were calculated by a two-sided one-sample t-test against a theoretical mean of 50%.

Alignment of these dominant transcripts revealed only five nucleotide mismatches, corresponding to >98% sequence identity (Fig. 1c,d). The near-identity of these transcripts complicates standard short-read RNA-seq quantification. To distinguish them, we leveraged diagnostic nucleotide differences between the dominant isoforms to assign reads unambiguously and quantify paralog-specific expression. Using this approach, we examined the relative abundance of these transcripts across several cell lines and consistently observed higher expression of CYTOR compared to MORRBID (Fig. 1e,f).

To determine whether these loci differ in local regulatory activity, we assessed the effect of transcriptional repression on neighboring genes. CRISPR interference (CRISPRi)-mediated repression of CYTOR and MORRBID had no significant effect on local PCGs within a 2-3 megabase window surrounding their transcription start sites, including the MORRBID head-to-head neighbor BCL2L11 (BIM) (Fig. 1g,h). These findings argue against locus-specific cis-regulatory functions and further support the conclusion that CYTOR and MORRBID operate through shared mechanisms despite their divergent genomic organization^12,22^. Together, these findings indicate that, despite their distinct genomic contexts, CYTOR and MORRBID produce nearly identical dominant transcripts and exhibit highly similar transcriptional and splicing patterns^13,23,24^.

### CYTOR and MORRBID are associated with cell adhesion and migration

Given that CYTOR and MORRBID produce dominant transcript isoforms that are nearly identical in sequence, we sought to determine whether they are associated with similar cellular contexts and biological processes. To this end, we first analyzed single-cell RNA sequencing data from CELLxGENE to identify the cell types in which these transcripts are most highly expressed. Across tissues, CYTOR expression was enriched in a consistent set of mesenchymal-associated cell types (supplementary Table1), including fibroblasts, stromal cells, vascular smooth muscle cells, and pericytes (Fig. 2a)^25–27^. Notably, these cell types overlapped with those expressing MORRBID (Supplementary Fig. S1a), consistent with the strong correlation observed between the two transcripts^26^.

**Fig. 2.**
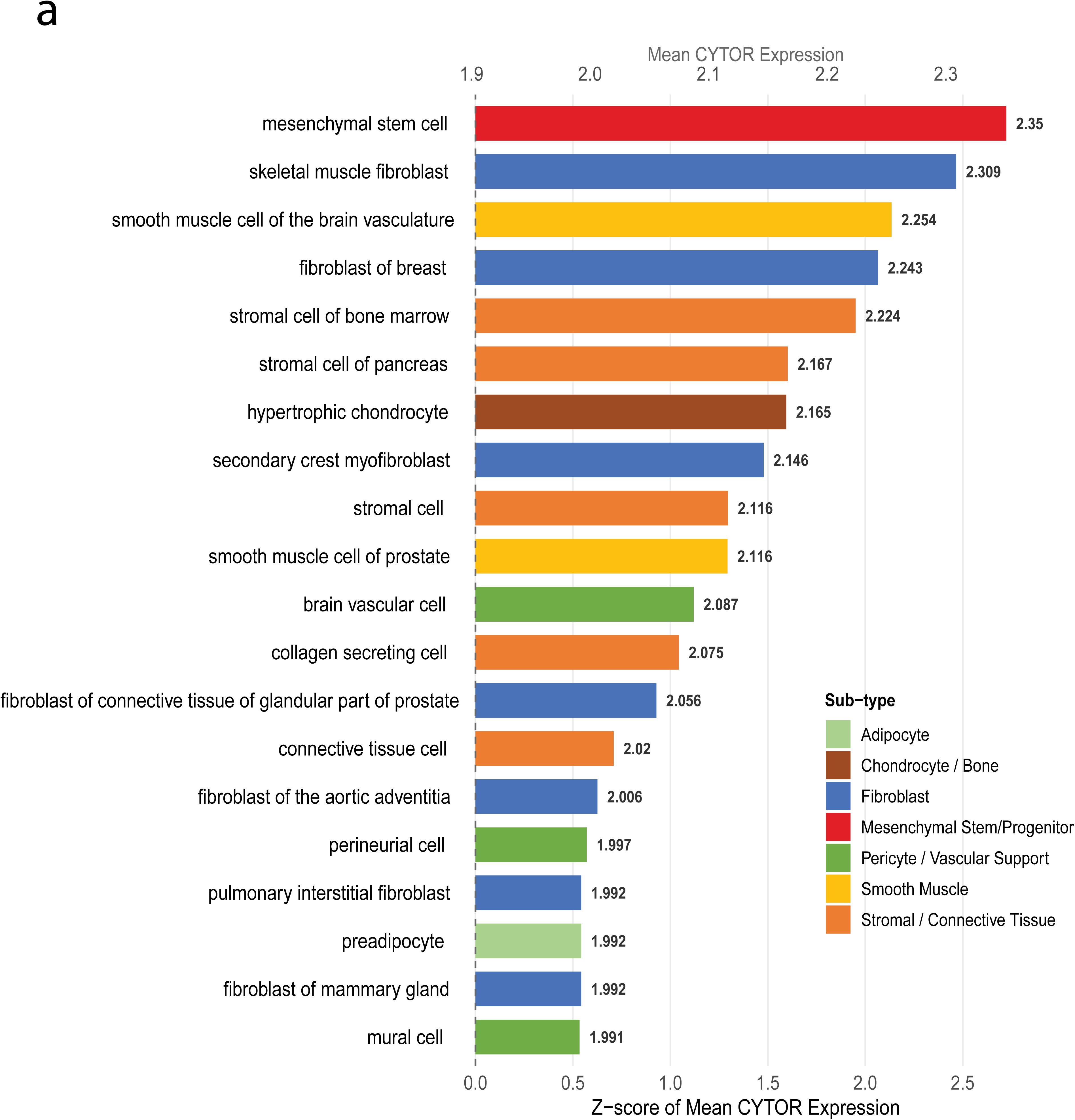
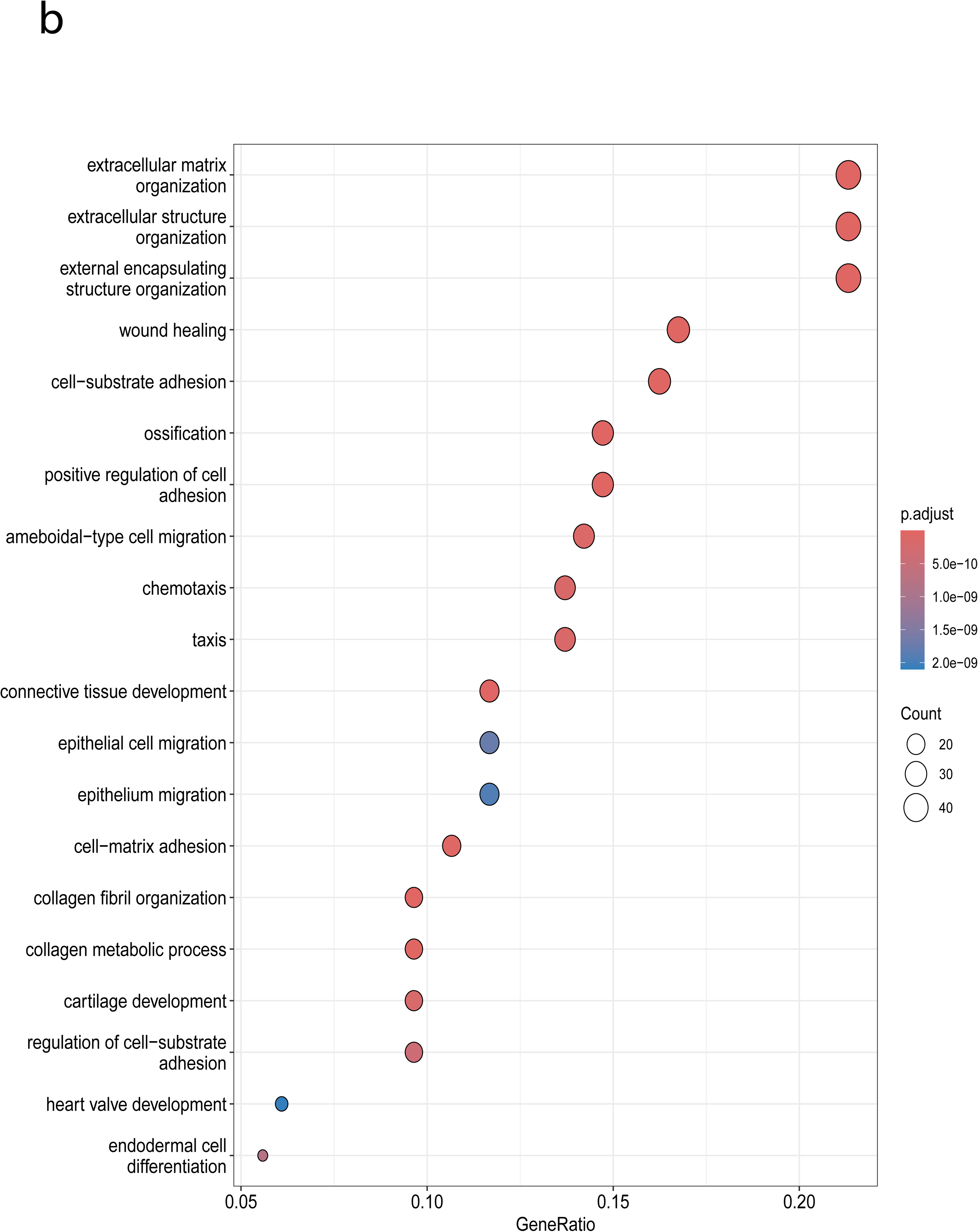
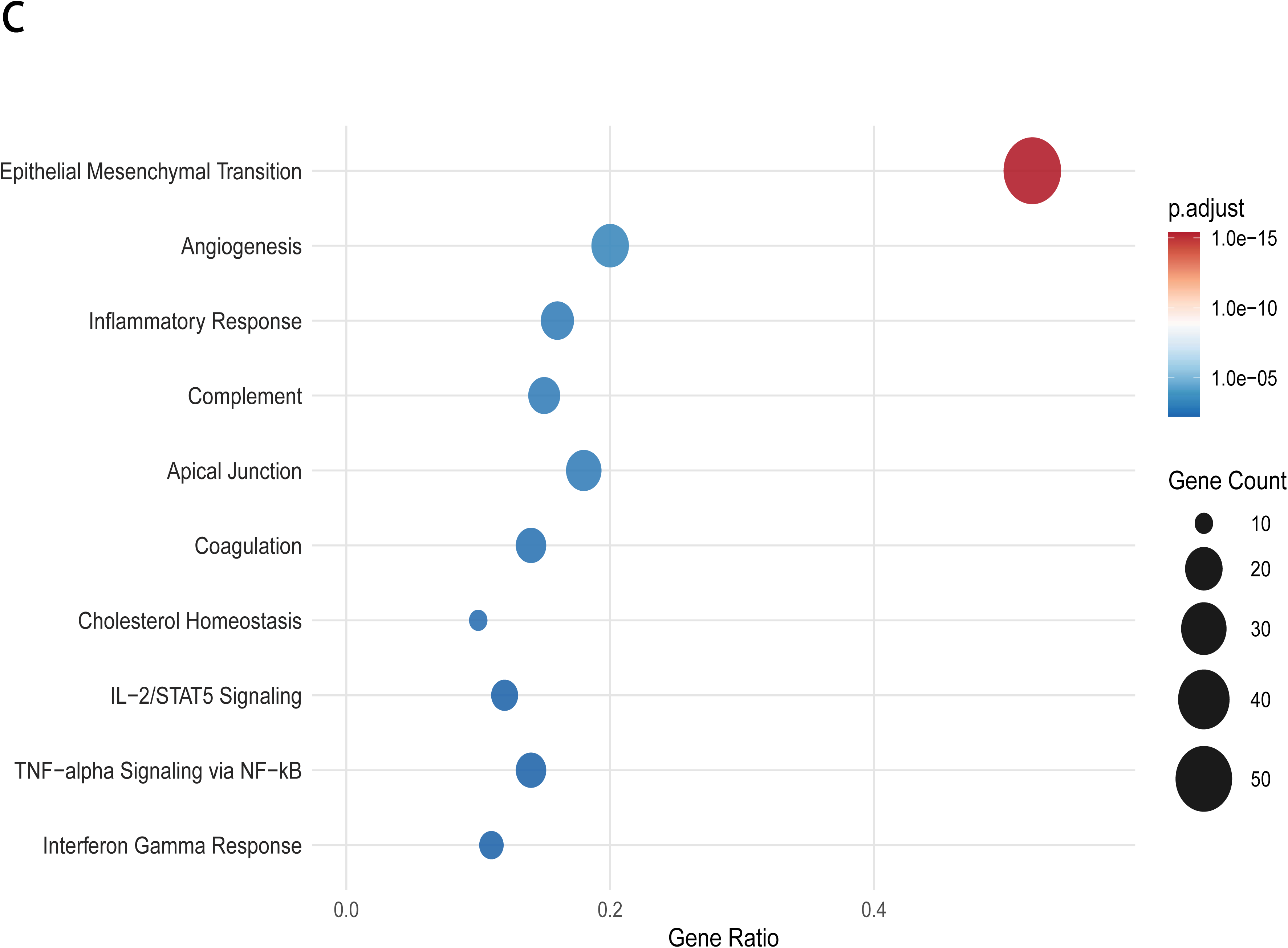
CYTOR and MORRBID are associated with mesenchymal cell identity and adhesion-related transcriptional programs. **a** Mean CYTOR expression across cell types from CELLxGENE single-cell RNA-sequencing (scRNA-seq) datasets, shown as Z-scores of mean expression across cell types. **b** Gene Ontology (GO) Biological Process enrichment analysis of genes co-expressed with CYTOR in GTEx bulk RNA-sequencing data.**c** Hallmark gene set enrichment analysis of the top CYTOR co-expressed genes across 32 TCGA cancer types.

To further characterize their functional associations, we identified hallmark gene sets for these enriched cell types and analyzed bulk RNA-sequencing (RNA-seq) data from the GTEx project. We computed gene expression correlations with CYTOR, and then performed Gene Ontology (GO) Biological analyses on the co-expressed genes (Fig. 2b) and for MORRBID (Supplementary Fig. S1b). This analysis revealed highly overlapping enrichment profiles, including pathways related to extracellular matrix organization, tissue migration including endothelia and epithelial, cell matrix adhesion process, and wound healing. These findings indicate that CYTOR and MORRBID are associated with a shared transcriptional program linked to cell adhesion, migration, and mesenchymal cell identity^23,26,28–30^. Consistent with this, CYTOR and MORRBID expression were highly correlated across non-diseased tissues (GTEx; r = 0.907), indicating coordinated regulation under physiological conditions (Supplementary Fig. S1c).

We therefore extended our analysis to cancer datasets, as both CYTOR and MORRBID are markedly upregulated across multiple cancer types, and their elevated expression is associated with tumor progression and poor clinical outcomes (Supplementary Fig.S1 d,e)^31,32^. We next examined the top 100 CYTOR co-expressed genes across 32 TCGA cancer types. Notably, CYTOR and MORRBID expression were highly correlated in tumor samples (TCGA; r = 0.873) (Supplementary Fig. S1f), further supporting shared regulation and function in cancer. Among the top co-expressed genes, 72 were consistently identified in more than five tumor types, then Gene set enrichment analysis revealed strong enrichment for the epithelial–mesenchymal transition (EMT) signature (Fig. 2c), consistent with a role in cell adhesion, migration, and tumor progression.

### CYTOR and MORRBID knockdown impairs cancer cell migration

To experimentally assess the cellular functions of CYTOR and MORRBID, we performed CRISPR interference (CRISPRi)-mediated repression in four human cancer cell lines representing distinct tumour types: MDA-MB-231 (triple-negative breast cancer), MCF7 (luminal ER-positive breast cancer), H1299 (non-small cell lung carcinoma), and U2OS (osteosarcoma). Because the CRISPRi target sequence is shared between the CYTOR and MORRBID promoters, this approach simultaneously represses both loci and therefore interrogates their combined function. Efficient knockdown (KD) across all four cell lines was confirmed by reverse transcription-quantitative PCR (RT-qPCR) (Fig. 3a). We then evaluated the phenotypic consequences of dual repression on major cancer-related processes, including cell cycle progression, apoptosis, and cell growth.

**Fig. 3.**
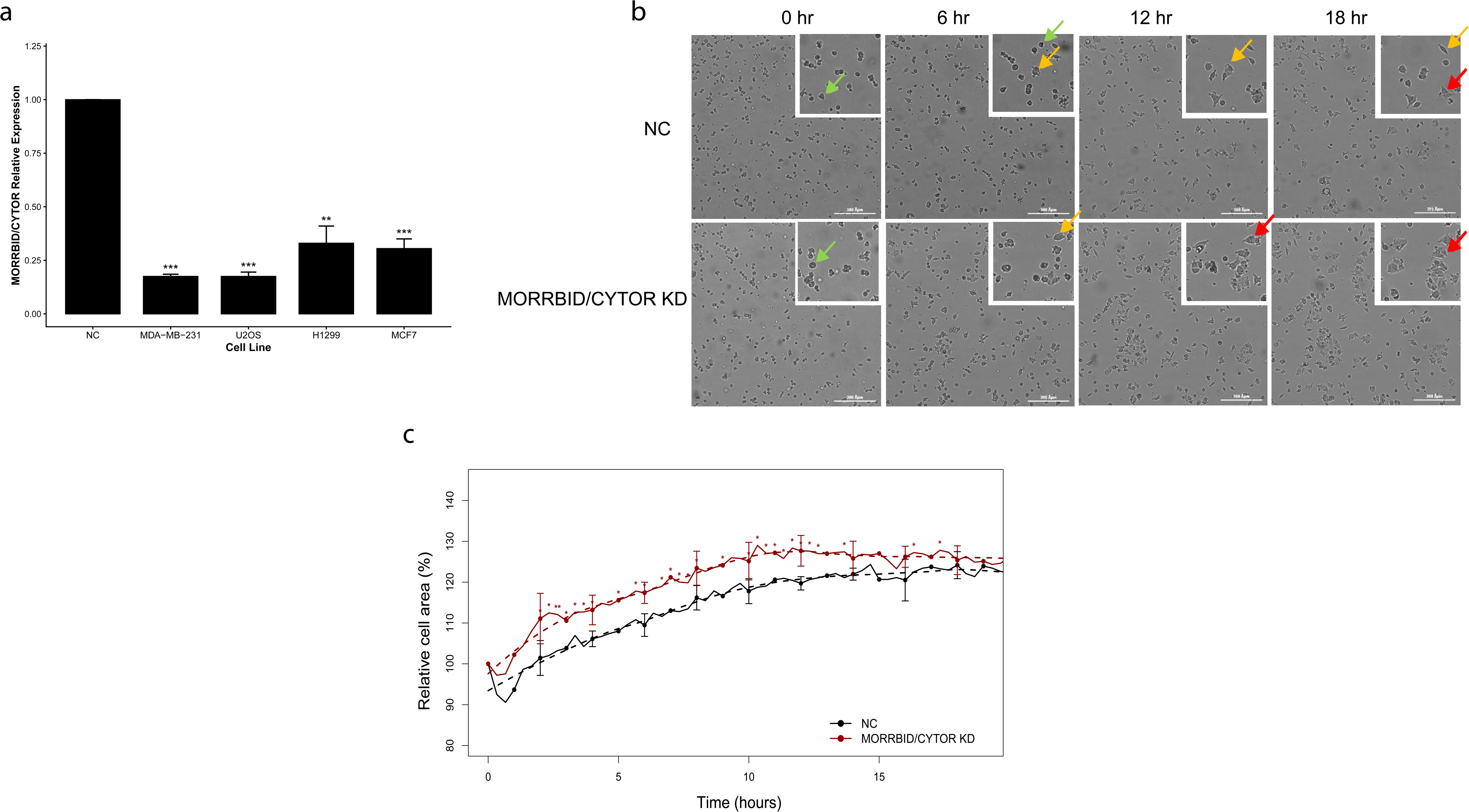
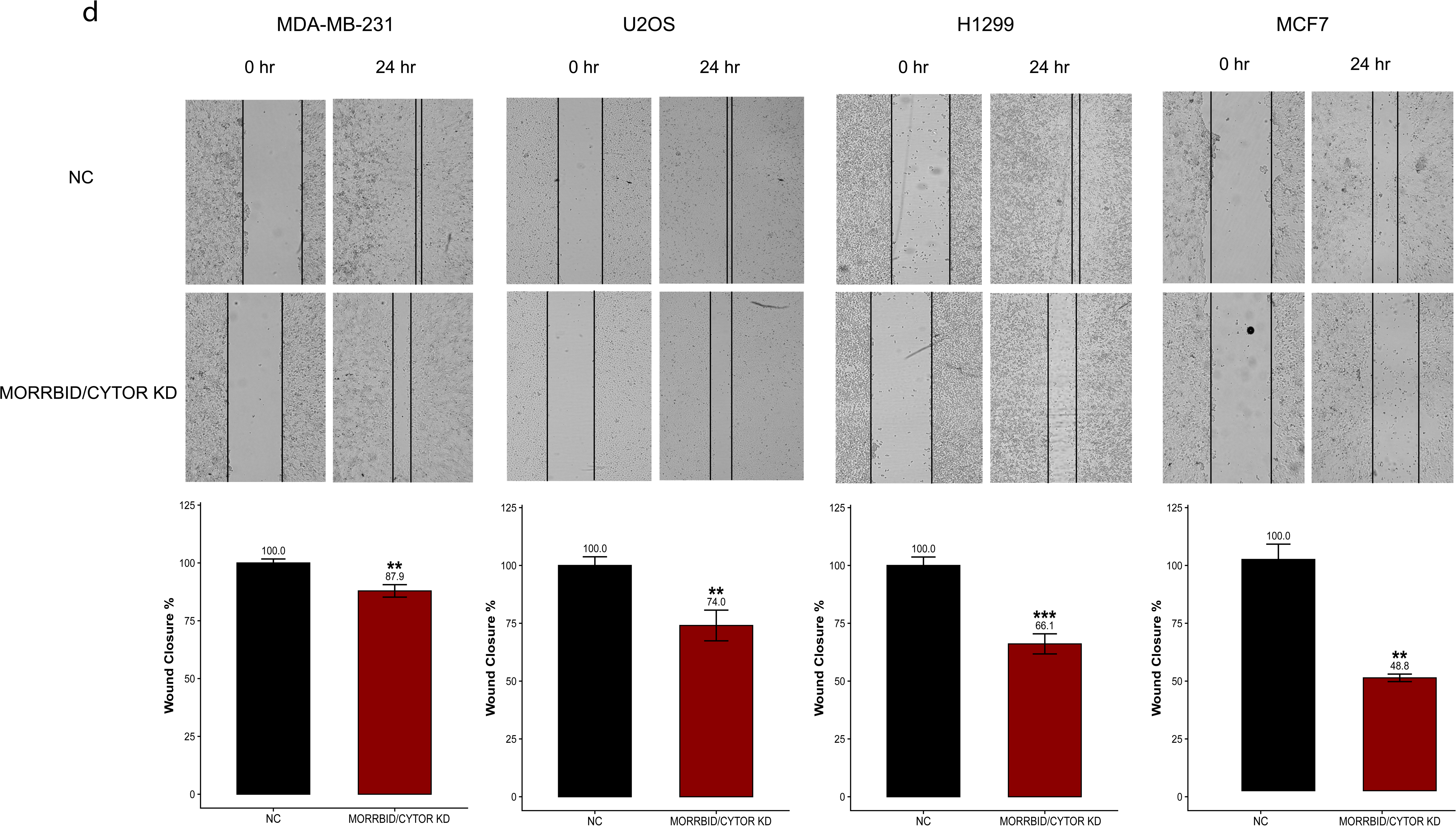
Knockdown of CYTOR and MORRBID increases cell attachment and impairs cancer cell migration. **a** Reverse transcription–quantitative polymerase chain reaction (RT–qPCR) validation of MORRBID/CYTOR knockdown (KD) in the MDA-MB-231, U2OS, H1299 and MCF7. Expression is shown relative to negative control (NC). Bars represent mean ±SD. n=3, * p value<0.05, ** p value<0.01 following student T-test. **b** Representative phase-contrast microscopy images of NC and MORRBID/CYTOR KD cells at selected time points, showing cell morphology and spreading. Green arrows indicate rounded suspended (non-adherent) cells; yellow arrows indicate cells initiating adhesion (early attachment/spreading); red arrows indicate fully adherent, with enlarged spread morphology. **c** Quantification of relative cell area (%) over 24 hours, normalized to the initial time point (t = 0). Data are presented as mean ± SD. n=3, * p value<0.05, ** p value<0.01 following student T-test. **d** Cell migration analysis by wound healing. Top. Brightfield microscopy footages of cells plate immediately following “scratch” and 24hr later. The black lines were added to represent the gap. Bottom, a normalized quantification of the gap at 24hr hours following the “scratch”. Bars represent mean ±SD. n>3, * p value<0.05, ** p value<0.01, *** p value <0.001. following student T-test.

In agreement with previous reports, simultaneous repression of CYTOR and MORRBID produced cell-type-specific phenotypes, with only moderate consistency and reproducibility across cell lines^33,34^. Specifically, cell cycle was not significantly affected in any of the tested models (Supplementary Fig. S2a). Apoptosis analysis revealed an increase in apoptotic cells only in MDA-MB-231 cells, with no comparable effect observed in H1299, U2OS or MCF7 cells (Supplementary Fig. S2b). Similarly, the impact of KD on cell growth was variable, with reduced proliferation in MDA-MB-231 cells, increased growth in H1299 cells, and no significant change in U2OS and MCF7 cells (Supplementary Fig. S2c).

However, closer examination after plating suggested that cells subjected to simultaneous repression of CYTOR and MORRBID exhibit altered adhesion dynamics. To assess this directly, we monitored cells after seeding by time-lapse microscopy in MDA-MB-231 cells and observed that simultaneous repression of CYTOR and MORRBID enhanced cell spreading, resulting in increased surface area and cell size compared with negative controls (NC), indicative of increased attachment to the culture surface (Fig. 3b,c and Supplementary Fig. 2d).

As cell spreading reflects the formation of cell-substrate adhesions and reorganization of the actin cytoskeleton ^35^, and given that excessive or stabilized adhesion can limit cell motility^36^, we further analyzed the effect of simultaneous repression of CYTOR and MORRBID on migration using a wound-healing assay. This revealed a consistent decrease in gap closure across four migratory cell lines (Fig. 3d)

To validate these findings using an independent approach, we performed KD of CYTOR and MORRBID using siRNAs that mainly targets cytoplasmic RNA. Migration was subsequently assessed using the same wound-healing assay. Notably, the results were reproducible and consistent with those obtained using the initial method, supporting that the observed effects are specific to CYTOR and MORRBID rather than attributable to the experimental system (Supplementary Fig.S3a-c).

Importantly, analysis of breast and lung cancer patient datasets revealed that elevated expression of CYTOR and MORRBID is associated with reduced overall survival (Supplementary Fig. S4a,b,d,e), highlighting their clinical relevance in promoting cell migration and mesenchymal programs. Consistently, both CYTOR and MORRBID are expressed at higher levels in advanced-stage tumors (stages III-IV), which are typically associated with metastatic disease (Supplementary Fig. S4c,f).

Furthermore, Analysis of publicly available datasets of breast and lung tissues, including both diseases and non-diseased samples, revealed robust enrichment of EMT pathways among genes co-expressed with CYTOR and MORRBID (supplementary Fig. S5a-d). This observation is consistent with our analyses across multiple tissues, reinforcing a role for CYTOR and MORRBID in mesenchymal gene regulation and cell migration, and demonstrating that this association is conserved across biological contexts rather than being restricted to cancer.

Although CYTOR and MORRBID have been linked to diverse cellular processes across different cancer types and experimental models, many of these reported functions appear to be context-dependent^21,34^. In contrast, our data indicate that simultaneous repression of CYTOR and MORRBID consistently affects cell migration across different cell lines, pointing to this phenotype as a shared and robust functional output.

Together, these findings indicate that CYTOR and MORRBID have a shared role in cancer cell adhesion and migration and provide a basis for subsequent mechanistic investigation.

### CYTOR and MORRBID promote cell migration in an EPHA4-dependent manner

To gain further insight into the mechanisms underlying this shared function of CYTOR and MORRBID, we performed transcriptomic analysis in the breast cancer cell line MDA-MB-231, a well-established model for studying cell migration and EMT^37^. RNA-seq following simultaneous repression of CYTOR and MORRBID identified 173 differentially expressed genes (Fig. 4a), most of which were downregulated. Consistent with the observed phenotype, gene set enrichment analysis identified significant enrichment of pathways related to cell migration, extracellular matrix organization, cell adhesion, and ERK signaling (Fig. 4b).

**Fig. 4.**
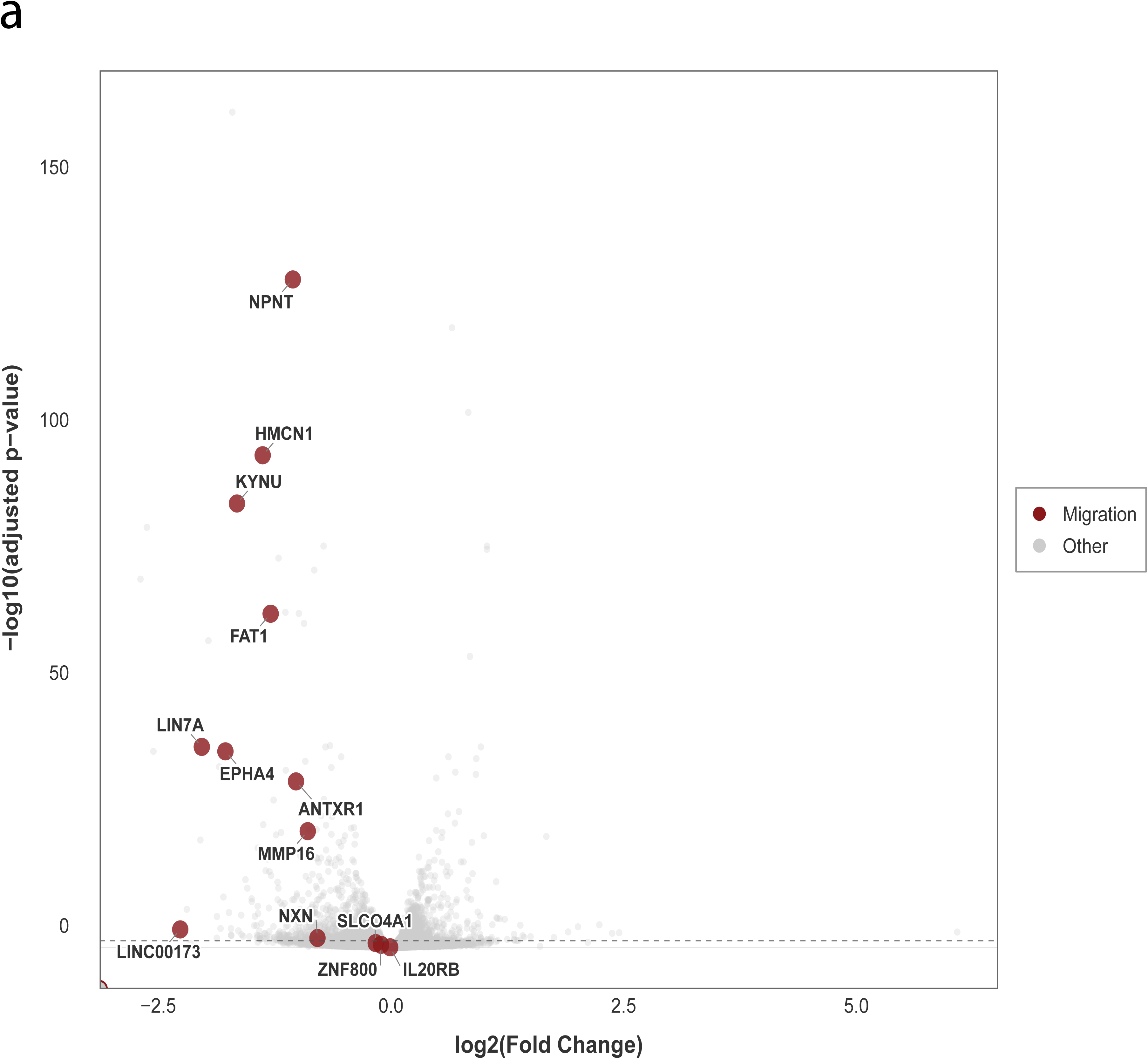
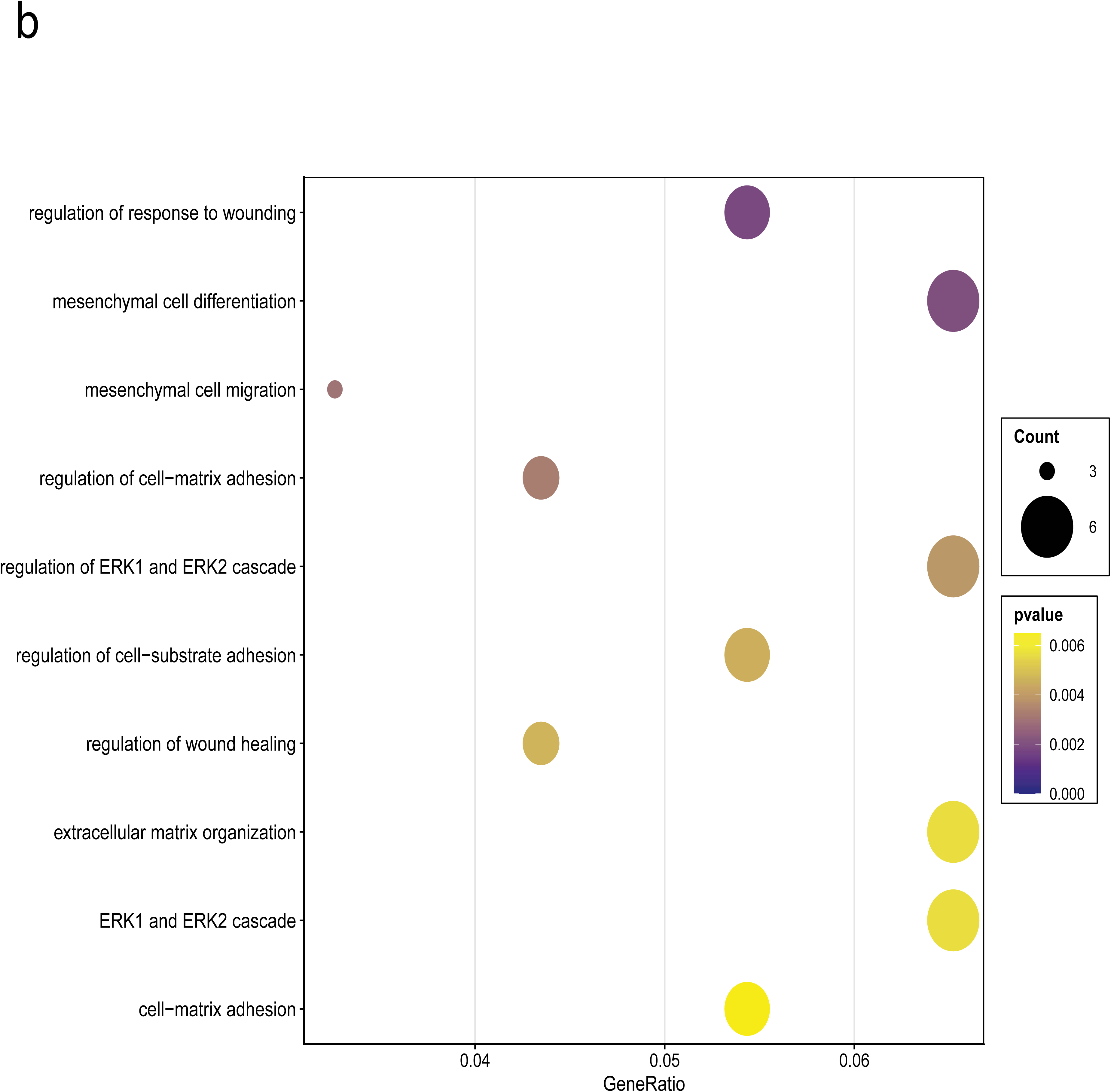
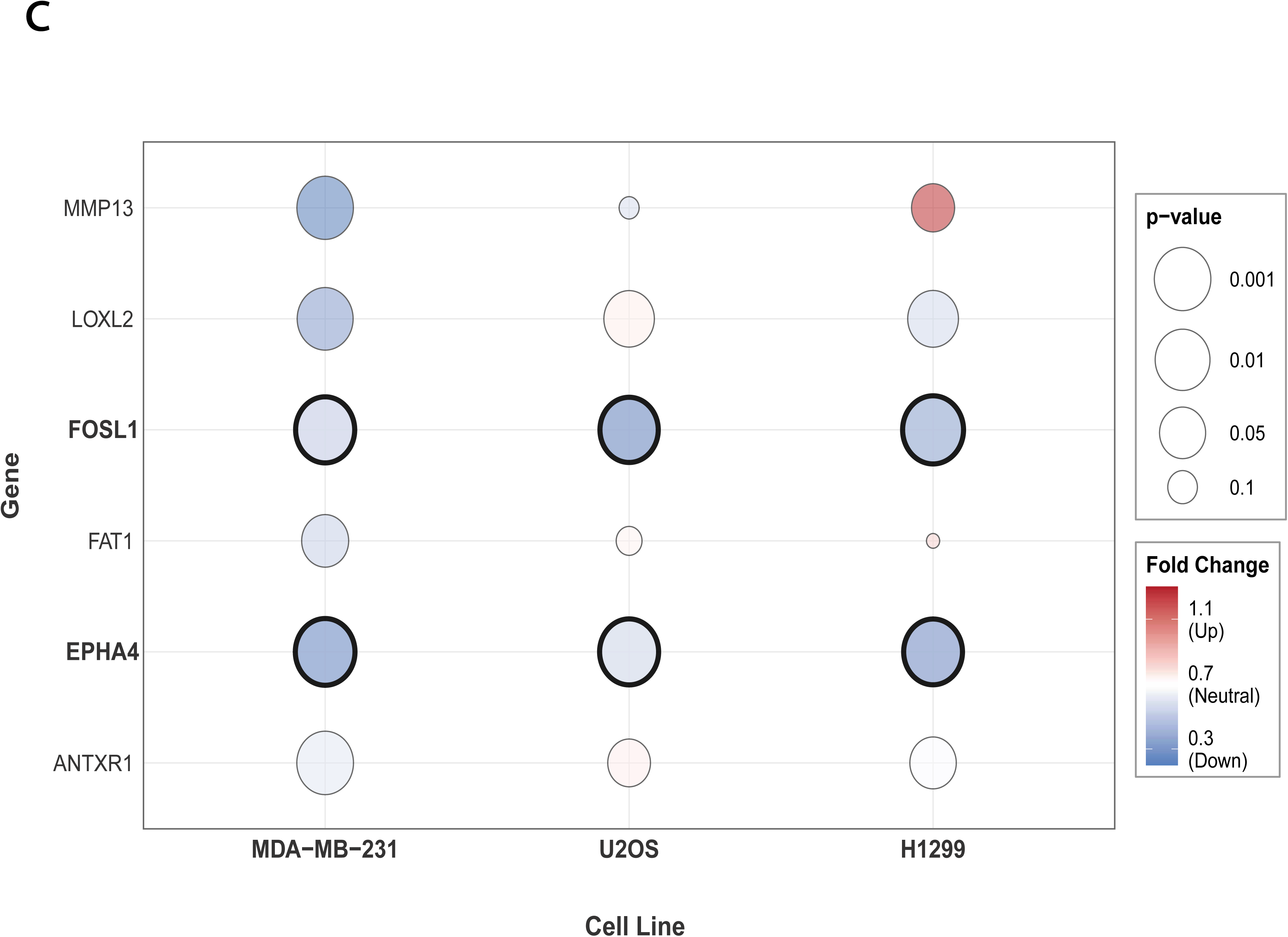
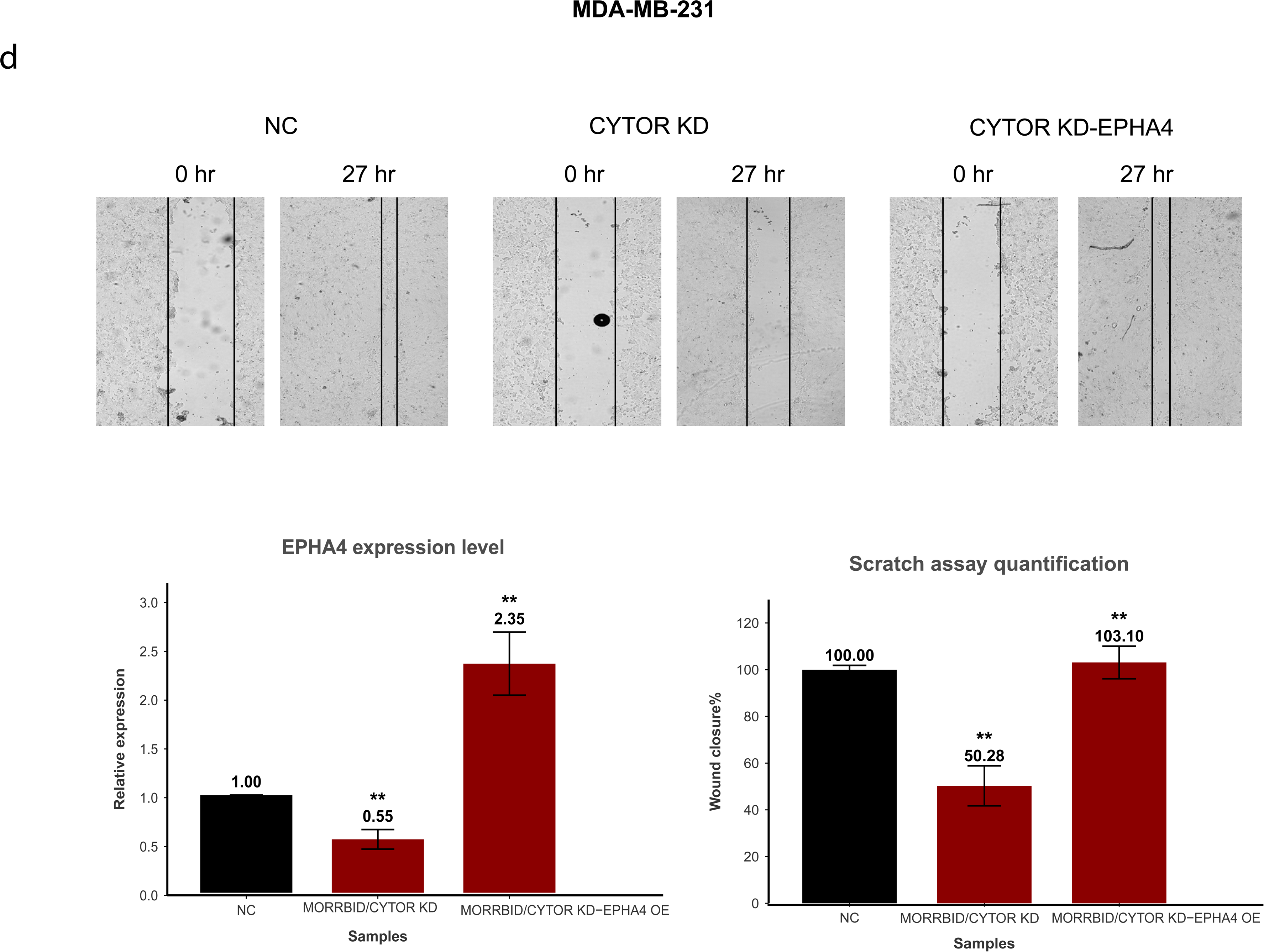
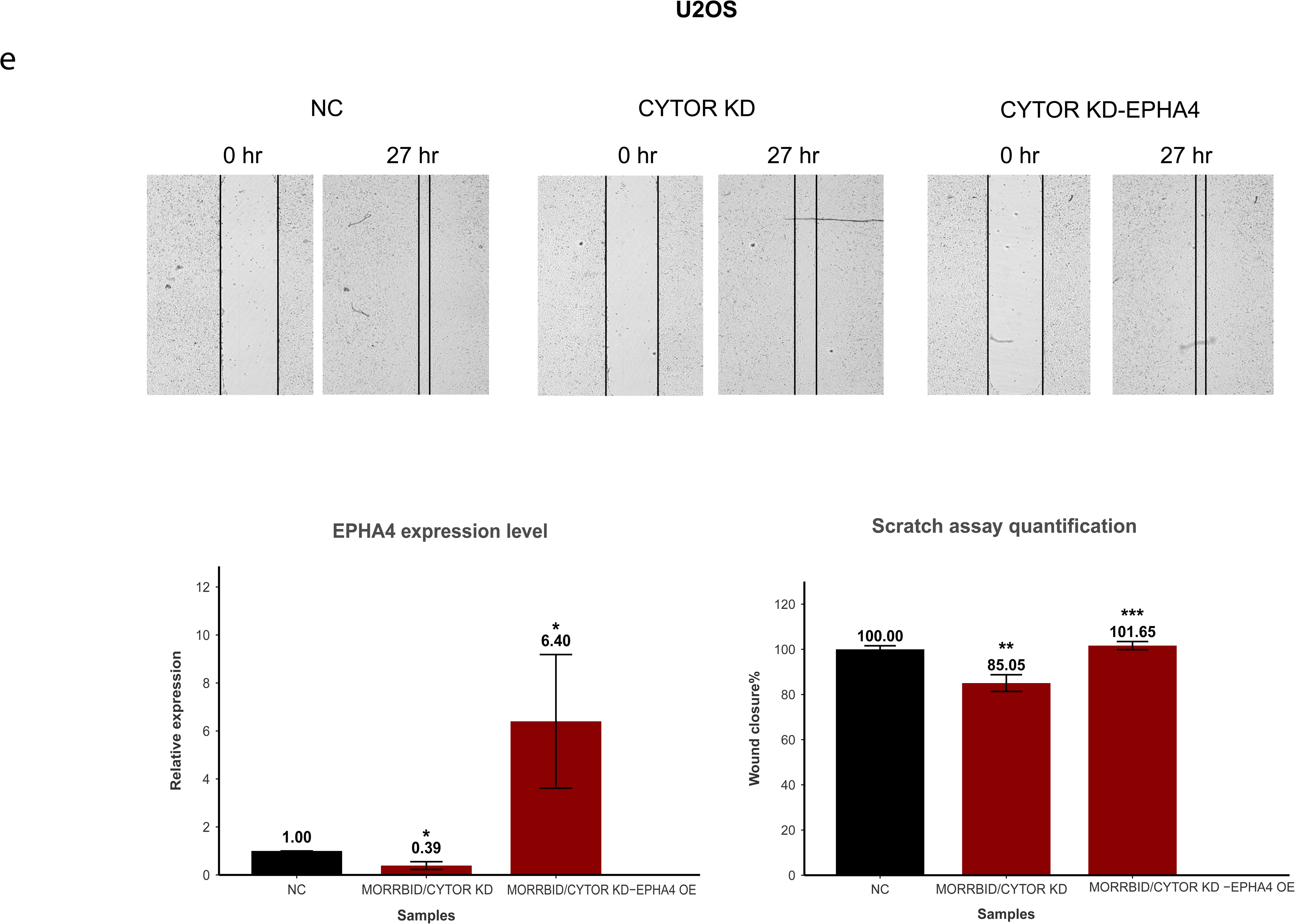
CYTOR and MORRBID promote cell migration through transcriptional regulation of EPHA4. **a** Volcano plot showing differentially expressed genes following simultaneous KD of MORRBID/CYTOR in MDA-MB-231 cells, as determined by RNA sequencing (RNA-seq). Migration-related genes are highlighted in red. **b** Pathway enrichment analysis of MORRBID/CYTOR-associated biological processes. **c** Quantitative polymerase chain reaction (qPCR) validation of selected migration genes following MORRBID/CYTOR KD in three cell lines. **d and e** Brightfield microscopy footages of cells plate immediately following “scratch” following EPHA4 overexpression, and 24hr later in MDA-MB-231 and U2OS. Bars represent mean ±SD. n=3, * p value<0.05, ** p value<0.01, *** p value <0.001.

To validate these findings, we selected a subset of migration-related genes for quantitative Polymerase Chain Reaction (qPCR) analysis based on prior literature (Supplementary Fig. S6a). Although several genes exhibited reduced expression following simultaneous repression of CYTOR and MORRBID in MDA-MB-231 cells, validation across additional cancer cell lines, including H1299 and U2OS, identified a consistent and significant reduction in the receptor tyrosine kinase ephrin type-A receptor 4 (EPHA4) across all cell types (Fig. 4c and Supplementary Fig. S6b). EPHA4 is an Eph family receptor tyrosine kinase that regulates cell-cell communication, adhesion, and migration through cytoskeletal remodeling. In tumor models, its effect on EMT are context-dependent, but EPHA4 has been linked to altered migratory and invasive behavior^38^. To test the role of EPHA4, we ectopically overexpressed it in cells subjected to simultaneous repression of CYTOR and MORRBID, which significantly rescued cell migration (Fig. 4d,e), indicating that EPHA4 functions as a key downstream effector of CYTOR- and MORRBID-mediated regulation of cell migration.

### CYTOR and MORRBID are localized in the cytoplasm and physically bind MEK2 and modulate MAPK signaling activity

To further investigate how CYTOR and MORRBID regulate cell migration at the molecular level, we first analyzed publicly available RNA antisense purification-mass spectrometry (RAP-MS) data generated in HeLa cells using oligonucleotide probes targeting CYTOR^13^. These data revealed enrichment of MEK2 among proteins associated with the lncRNA transcripts. MEK2 (MAP2K2) is a predominantly cytoplasmic dual-specificity kinase and a central component of the RAS-RAF-MEK-ERK signaling cascade. By phosphorylating and activating ERK1/2, MEK2 controls pathway output and thereby regulates fundamental cellular processes, including proliferation, survival, differentiation, and migration^39^. Although MEK2 is not considered a canonical RNA-binding protein, its association with lncRNAs may occur through non-canonical RNA-binding interfaces or indirect recruitment within ribonucleoprotein complexes, as has been increasingly recognized for many cytoplasmic signaling proteins lacking classical RNA-binding domains^40^. This interpretation is consistent with recent evidence showing that lncRNAs can directly regulate the localization and function of non-canonical RNA-binding signaling factors^40^. Importantly, the RAP-MS data provide a plausible mechanistic link to our transcriptomic analysis, which showed enrichment of differentially expressed genes associated with ERK signaling.

Because MEK2 is predominantly localized in the cytoplasm, we next examined the subcellular distribution of CYTOR and MORRBID. Single-molecule RNA fluorescence in situ hybridization (smRNA-FISH) confirmed that these lncRNAs are enriched in the cytoplasmic compartment (Fig. 5a and supplementary Fig. S7).

**Fig. 5.**
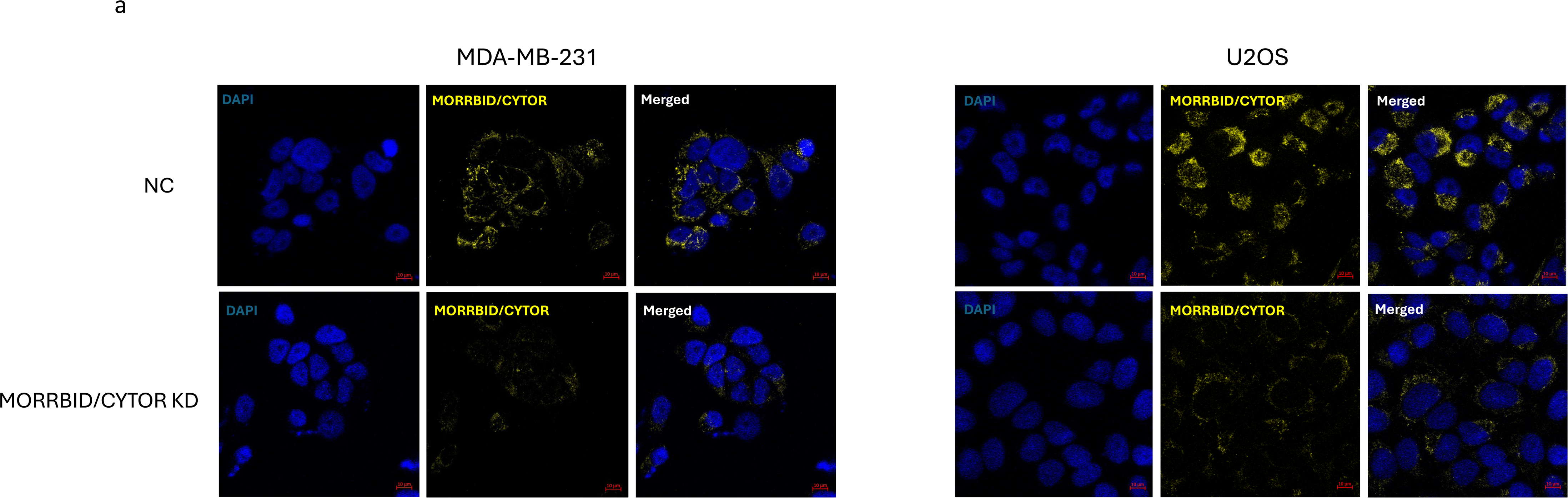
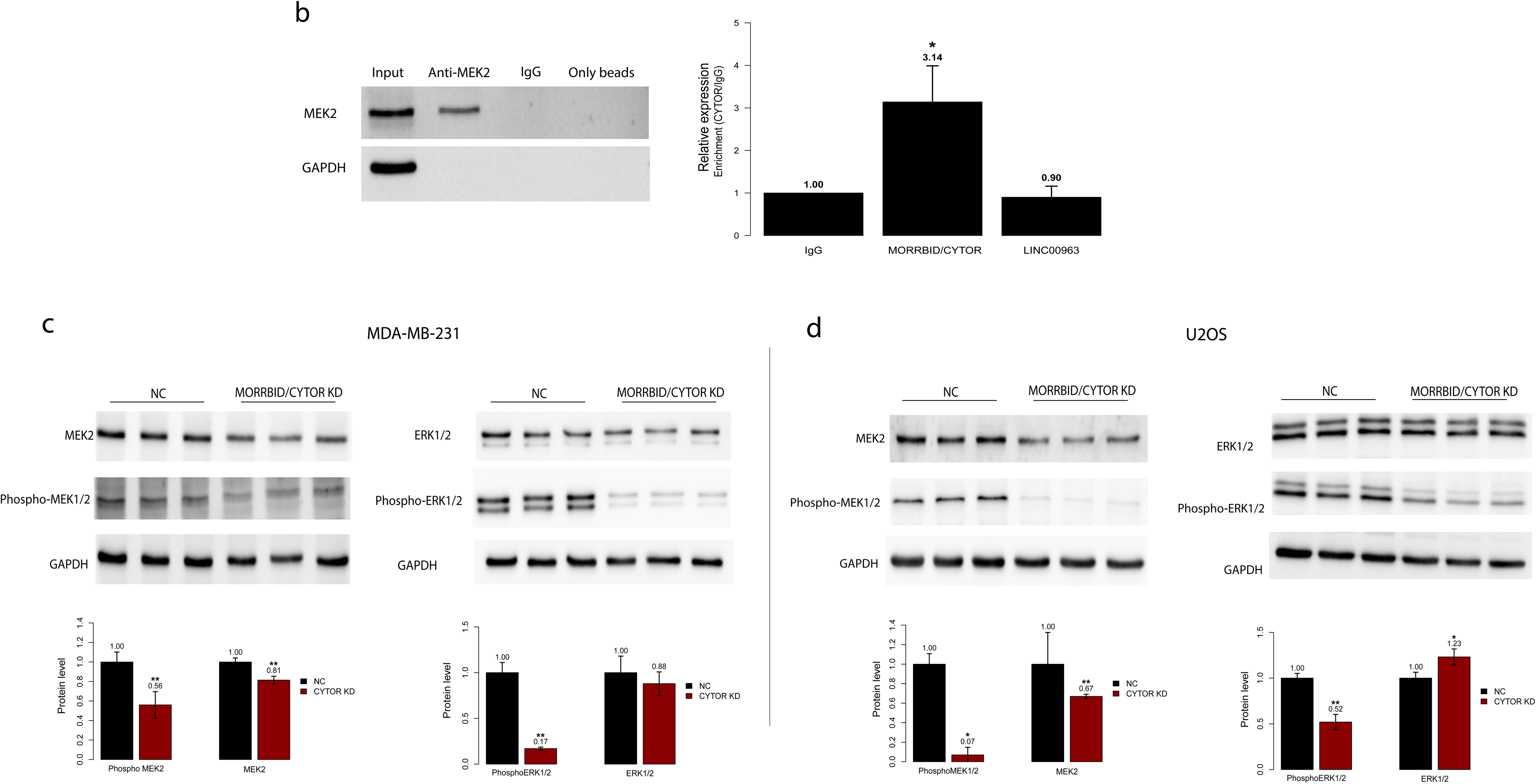
CYTOR and MORRBID are cytoplasmic lncRNAs that associate with MEK2 and regulate MAPK signalling activity. **a** Representative RNA in situ hybridization images showing CYTOR localization in MDA-MB-231 cells (left) and U2OS cells (right).n=3 biological repeats. **b** Cross-linked immunoprecipitation (CLIP) of endogenous MEK2 in MDA-MB-231 cells followed by qPCR analysis demonstrating interaction between CYTOR/MORRBID transcripts and MEK2. Bars represent mean ±SD. n=3, * p value<0.05, ** p value<0.01. **c** and **d** Representative Western blot (WB) analysis and quantification of MEK1/2, phospho-MEK1/2 (p-MEK1/2), ERK1/2, and phospho-ERK1/2 (p-ERK1/2) following MORRBID/CYTOR KD in MDA-MB-231 and U2OS cells. n>3, * p value<0.05, ** p value<0.01 following student T-test.

We next assessed whether MEK2 associates with the shared CYTOR/MORRBID transcript pool. Cross-linking immunoprecipitation followed by qPCR (CLIP-qPCR) demonstrated significant enrichment of CYTOR/MORRBID transcripts in MEK2 immunoprecipitates relative to both the IgG negative control and the cytoplasm-enriched lncRNA LINC00963^41^, which served as a negative control for MEK2 binding (Fig. 5b). These findings support a selective physical association between MEK2 and the shared CYTOR/MORRBID transcript pool.

To determine the functional consequences of the CYTOR/MORRBID-MEK2 interaction, we examined pathway activity following simultaneous repression of CYTOR and MORRBID. Notably, in both MDA-MB-231 and U2OS cells, we observed a significant but modest reduction in MEK2 protein levels. In contrast, MEK phosphorylation was reduced much more strongly (Fig. 5c,d). Consistent with the role of MEK2 as an upstream regulator of ERK, we also observed a strong reduction in ERK phosphorylation (Fig. 5c,d), indicating that CYTOR and MORRBID modulate signaling activity primarily through regulation of kinase activation rather than protein abundance.

Together, these findings identify CYTOR and MORRBID as cytoplasmic lncRNAs that physically associate with MEK2 and promote MEK-ERK signaling activity, supporting a role for these lncRNAs in regulating signaling pathways linked to cell migration.

### ERK-dependent AP-1 transcription factors link MAPK signaling to EPHA4 expression

Following phosphorylation by MEK, ERK becomes activated and rapidly translocates to the nucleus, where it regulates transcription factors (TFs) and contributes to chromatin remodeling and transcriptional control^39^. Using the GeneHancer database^42^, we identified four major regulatory elements associated with EPHA4; one located at the promoter and three within the gene body that interact with the promoter region (Fig. 6a). All four elements, as well as their promoter interactions, were classified as high-confidence “double elite” associations, indicating strong support for both the regulatory element itself and its interaction with the target promoter (Fig. 6a).

**Fig. 6.**
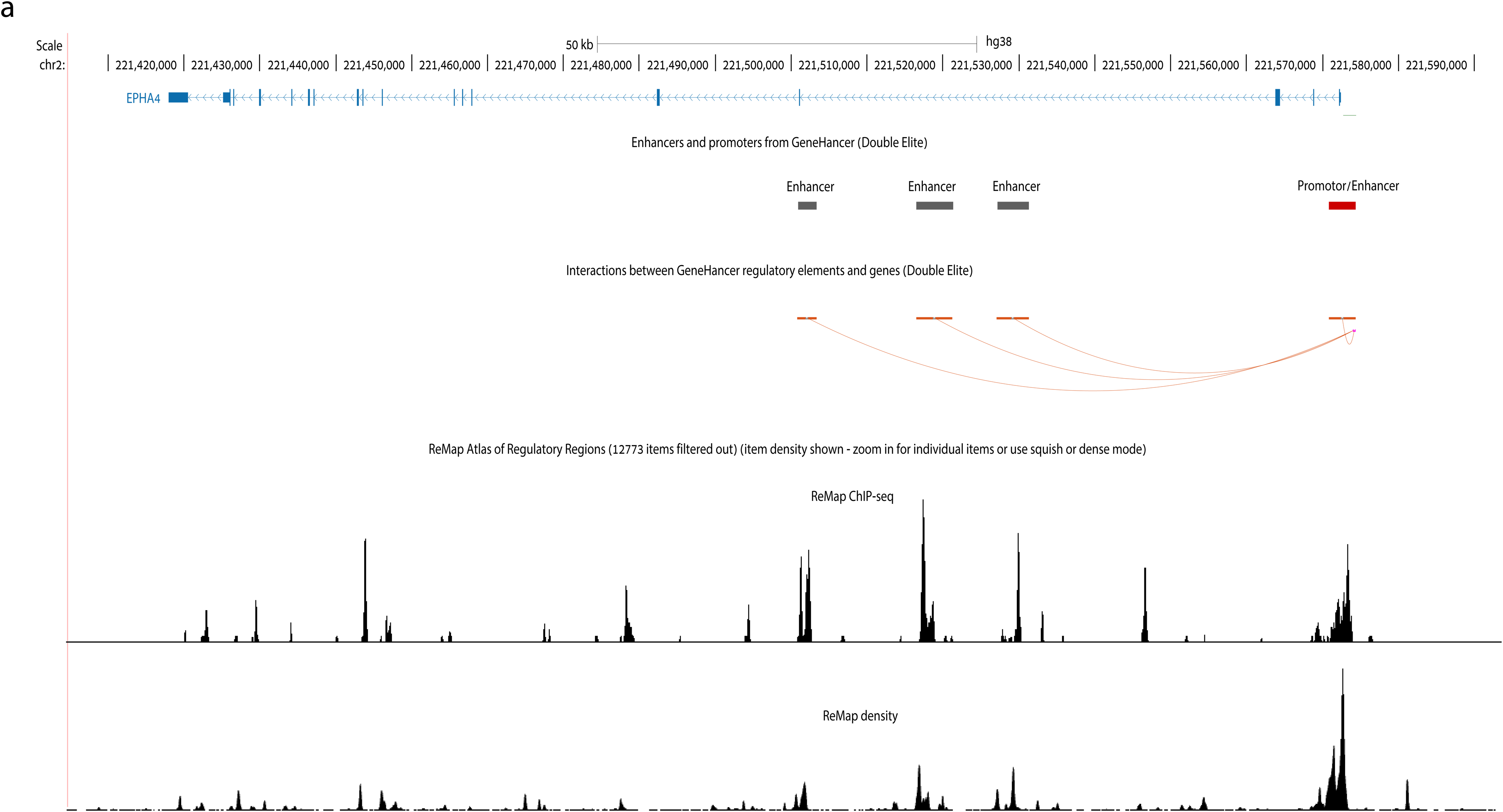
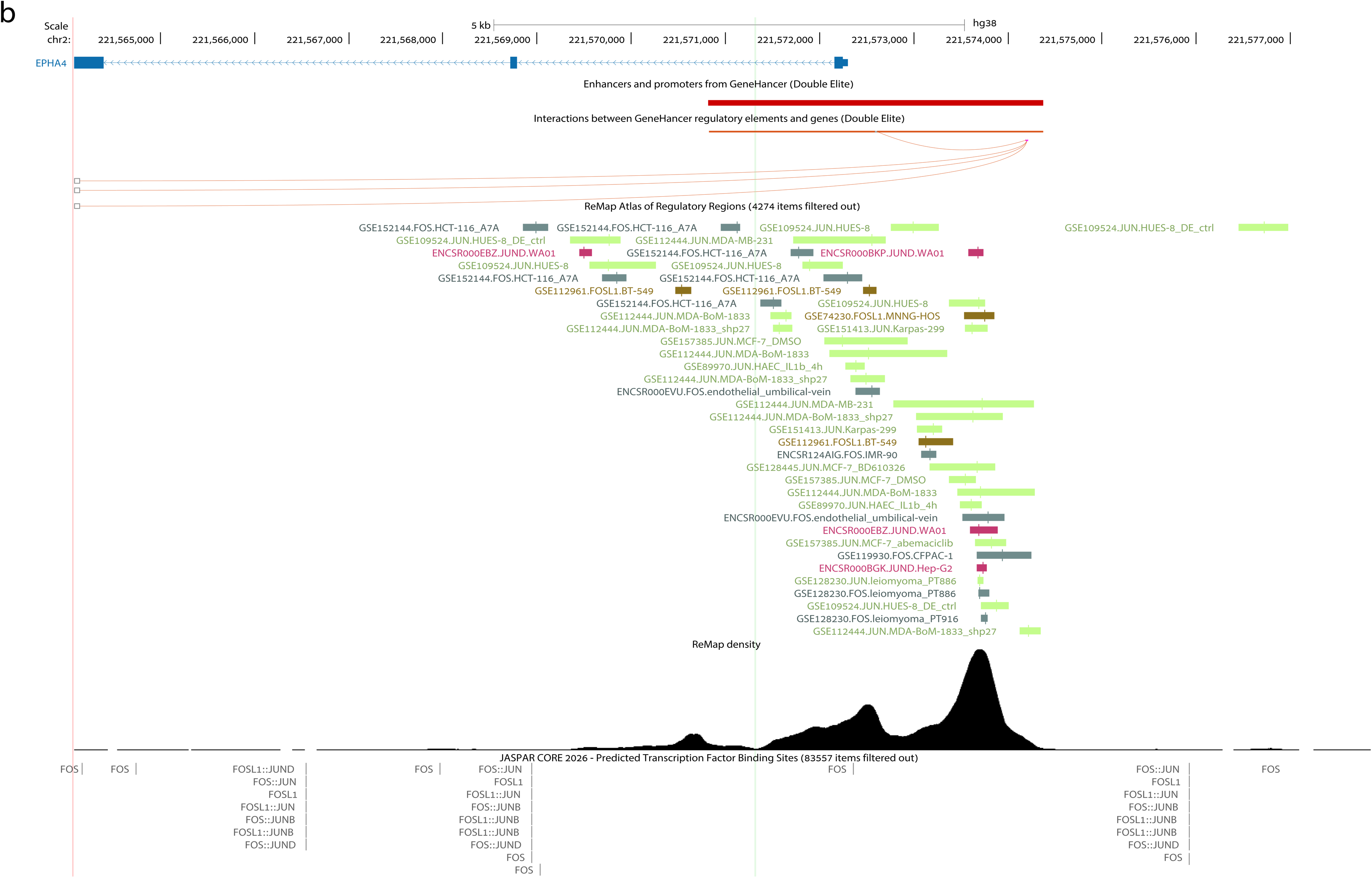
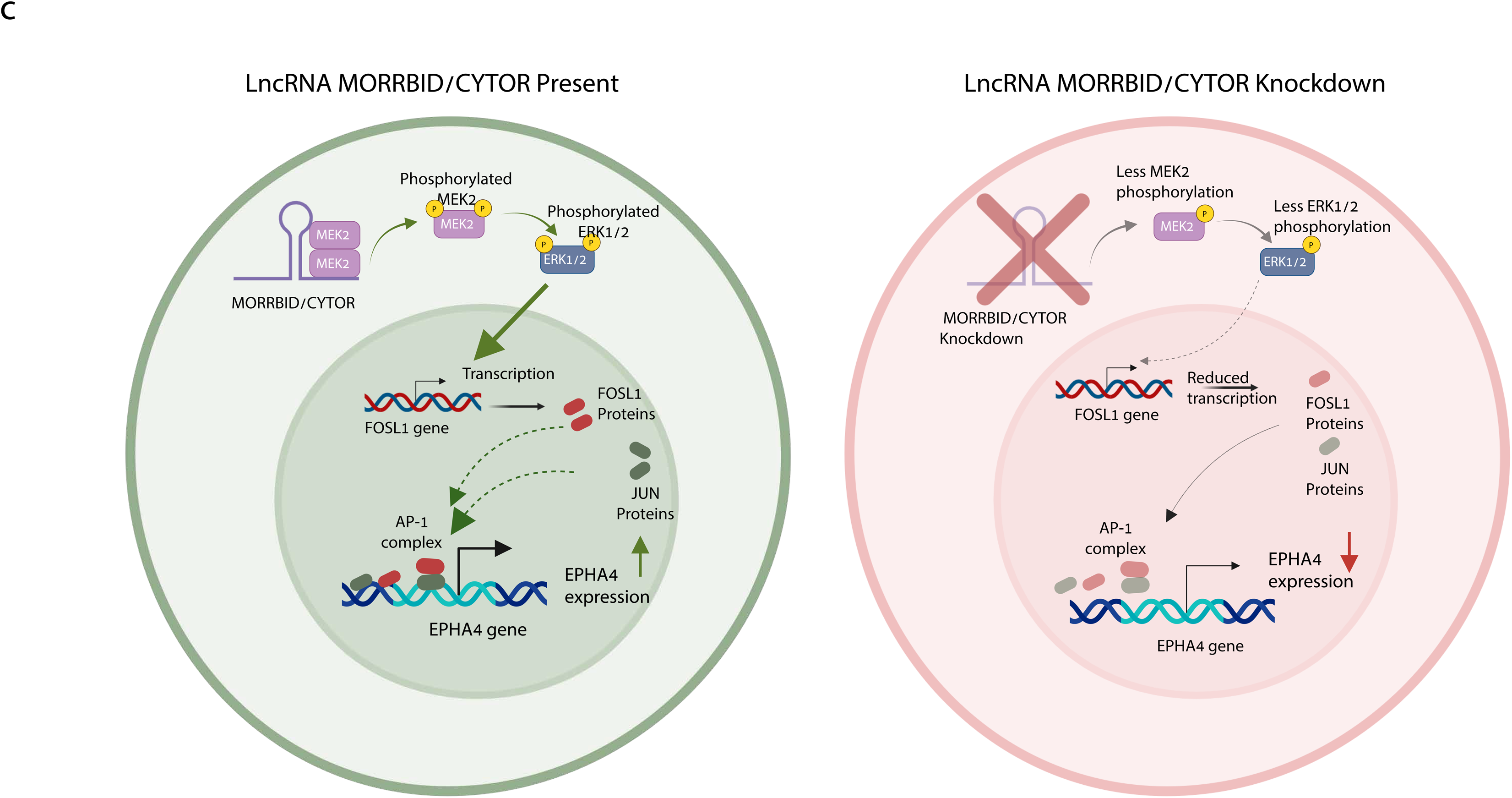
ERK-dependent AP-1 transcription factors are enriched at EPHA4 regulatory elements. **a** Schematic overview of the EPHA4 locus showing four major GeneHancer regulatory elements, including one promoter-associated element and three intragenic enhancers. All four elements display high-confidence “double elite” enhancer–promoter interactions with the EPHA4 promoter region. ReMap ChIP–seq density tracks indicate enrichment of transcription factor binding across these regulatory regions. **b** Zoomed view of the EPHA4 promoter-associated regulatory region showing ReMap ChIP–seq binding sites for JUN-, JUND-, FOS-, and FOSL1-family transcription factors, together with predicted AP-1-related motifs identified by JASPAR. Strong enrichment of FOSL1::JUN, FOSL1::JUND, FOS::JUN, FOS::JUNB, and FOS::JUND binding motifs is observed within the EPHA4 promoter region. **c** Proposed model for CYTOR- and MORRBID-mediated regulation of EPHA4 expression through the MEK2–ERK–AP-1 signaling axis. In the presence of CYTOR and MORRBID, interaction with MEK2 promotes MEK2 and ERK1/2 phosphorylation, resulting in increased FOSL1 expression and AP-1 complex activity, thereby enhancing EPHA4 transcription. Upon CYTOR and MORRBID knockdown, reduced MEK2 and ERK1/2 phosphorylation leads to decreased FOSL1 expression, impaired AP-1-mediated transcription, and reduced EPHA4 expression. Created in BioRender. Bester, A. (2026) https://BioRender.com/5b9a08f

To investigate the transcriptional regulators acting at these regions, we analyzed publicly available ChIP–seq datasets using ReMap^43^. This analysis identified binding sites for JUN- and FOS-family transcription factors within EPHA4 regulatory regions, suggesting that these factors may directly regulate EPHA4 transcription (Fig. 6b and Supplementary Fig. S8a-c). These transcription factors are well-established downstream effectors of ERK signaling^44,45^. Among the enriched factors, FOSL1 was particularly prominent (Fig. 6b), further supporting a role for CYTOR and MORRBID in MAPK pathway activity.

Consistent with this model, simultaneous KD of CYTOR and MORRBID led to reduced FOSL1 expression across three cell lines (Supplementary Fig. S6c). FOSL1 (FRA-1), a member of the FOS family and a well-characterized downstream effector of ERK signaling, is regulated by ERK at multiple levels^46^. At the transcriptional level, ERK activates ELK1 and c-JUN, which bind TPA response element (TRE) and Serum Response Element (SRE) elements in the FOSL1 promoter to induce transcription^47,48^. At the post-translational level, ERK directly phosphorylates FOSL1 at Ser252 and Ser265, thereby stabilizing the protein and protecting it from proteasomal degradation^49–51^.

FOSL1 forms heterodimeric complexes with JUN-family proteins as part of the AP-1 transcription factor complex, functionally analogous to canonical FOS–JUN dimers^45,52^. To further investigate this regulatory relationship, we performed motif scanning using JASPAR ^53^, and identified predicted binding sites for FOSL1::JUN heterodimers, as well as additional AP-1-related transcription factor complexes, within EPHA4 regulatory regions (Fig. 6b and Supplementary Fig. S8a-c).

Together, these findings suggest that CYTOR and MORRBID regulate EPHA4 expression through modulation of ERK-dependent AP-1 transcriptional activity. In this model, reduced ERK signaling following CYTOR and MORRBID depletion diminishes FOSL1 expression and activity, thereby impairing AP-1-mediated transcriptional activation of EPHA4.

### Model for CYTOR–MEK2–ERK regulation of EPHA4 expression and cancer cell migration

Based on our findings, we propose a model in which CYTOR and MORRBID function in the cytoplasm to regulate signaling activity through physical association with MEK2, thereby promoting activation of the MEK–ERK pathway. Enhanced ERK signaling, in turn, drives the expression of migration-associated genes, including EPHA4, likely through ERK-responsive transcriptional programs involving AP-1 transcription factors such as FOSL1. Activated ERK promotes FOSL1 expression, which may subsequently contribute to AP-1-mediated activation of EPHA4 transcription.

Conversely, depletion of CYTOR and MORRBID disrupts MEK2 and ERK1/2 phosphorylation, resulting in reduced FOSL1 expression, decreased EPHA4 levels and, consequently, impaired cancer cell migratory capacity (Fig. 6c).

Together, our results establish CYTOR and MORRBID as functionally indistinguishable cytoplasmic lncRNAs that promote cell adhesion and migration through a shared MEK2–ERK–AP-1–EPHA4 signaling axis. These findings unify previously fragmented and often model-specific observations, demonstrating that despite their distinct genomic contexts, these paralogous lncRNAs converge on a common molecular program. More broadly, our study refines the current view of CYTOR and MORRBID function and provides a mechanistic example of how duplicated lncRNAs can retain a shared functional output in cancer.

## Discussion

A defining feature of lncRNAs is their high evolutionary turnover. Many emerge rapidly through mechanisms such as segmental duplication, transposable element insertion, and local sequence change^54,55^. At the same time, lncRNA loci often evolve under relatively weak selective constraint^56^, raising the question of how newly emerged transcripts acquire biologically meaningful function. Once established, lncRNA expression and activity are further shaped by genomic context, which can enable the emergence of lineage-specific regulatory properties^57^. The CYTOR-MORRBID paralog pair provides a stringent test of this problem because CYTOR arose through primate-specific duplication and relocation of an ancestral MORRBID-like locus. Remarkably, our results show that this event did not generate two fundamentally different lncRNAs. Instead, CYTOR and MORRBID remain functionally inseparable, preserving a shared cytoplasmic MEK-ERK-dependent program that drives adhesion and migration. More broadly, these findings argue that lncRNA evolution can preserve biological output despite major changes in genomic context.

This conclusion directly challenges the prevailing view that MORRBID and CYTOR evolved into mechanistically distinct lncRNAs. MORRBID has been primarily framed as a nuclear, cis-acting regulator^12,31^, whereas CYTOR has largely been studied as a cytoplasmic promigratory lncRNA in cancer^58^, and both loci were subsequently linked to additional context-dependent functions through distinct proposed mechanisms^22,59^. That interpretation is further reinforced by transcript complexity at both loci: each produces multiple annotated isoforms, including transcripts that are nearly entirely shared across the duplicated region and others that contain locus-specific sequence. Our findings shift attention to the dominant shared two-exon transcript and argue that the clearest and most reproducible function of this paralog pair is a common cytoplasmic program linked to adhesion, migration, and MAPK signaling. At the same time, the presence of additional isoforms leaves open the important possibility that, under particular cellular contexts or conditions, locus-specific transcripts contribute more strongly and support additional CYTOR- or MORRBID-specific functions. In this view, transcript diversification after duplication may have added regulatory specialization without displacing a conserved core functional output.

This shared functional output is further supported by a convergent mechanistic framework. The physical association between MEK2 and the CYTOR/MORRBID transcript pool, together with the requirement for CYTOR and MORRBID in maintaining MEK and ERK phosphorylation, places these lncRNAs within a defined signaling pathway. Importantly, the MEK-ERK axis may account for several phenotypes previously attributed to CYTOR and MORRBID, including effects on migration, apoptosis, and cell-cycle regulation, and therefore provides an important complement to models centered on miRNA sponging ^12,13,22,27,60,61^. Although the precise nature of the RNA-protein interaction remains unresolved, our findings are consistent with the emerging role of non-canonical RNA-binding proteins in RNA-mediated regulation ^40^, and establish a unifying mechanistic framework for the shared function of this paralog pair.

In cancer, CYTOR and MORRBID define a shared functional unit that contributes to tumor progression by sustaining MAPK-ERK signaling and thereby promoting adhesion- and migration-associated programs. Within this framework, EPHA4 emerges as a consistent downstream effector of CYTOR/MORRBID-dependent ERK activity and a plausible mediator of the migratory phenotype. Mechanistically, this regulation may be mediated through ERK-responsive AP-1 transcription factors, particularly FOSL1, which was enriched at EPHA4 regulatory regions and downregulated following CYTOR/MORRBID depletion.

EPHA4 is a context-dependent Eph receptor tyrosine kinase that regulates cell adhesion, cytoskeletal remodeling, and migration, with reported roles in either promoting or constraining invasion depending on tumor context. Consistent with this interpretation, EPHA4 has also been implicated more broadly in processes involving tissue remodeling and cell migration, including neuronal development, wound healing, fibrosis, and immune cell movement ^38^. Although our data highlight EPHA4 as one important downstream effector of cancer cell migration, the CYTOR/MORRBID-MEK-ERK axis is likely to regulate additional targets in a context-dependent manner. Such broader downstream signaling may help explain the wide range of phenotypes previously associated with CYTOR and MORRBID.

An important implication of these findings is that the CYTOR/MORRBID axis should not be viewed solely through the lens of cancer. CYTOR has already been implicated in several nonmalignant settings, including exercise-responsive myogenesis and maintenance of fast-twitch muscle identity^62^, and delivery of a chemically modified CYTOR exon 2 domain has been shown to improve dystrophic myotube function^63^, T-cell activation and HIV gene regulation ^64^, and mechanosensitive immunomodulatory programs in mesenchymal stromal cells ^26^. On the MORRBID side, normal physiological roles have been described in short-lived myeloid cells^12^, and MIR4435-2HG has also been linked to metabolic function in primary myeloid dendritic cells ^65^. In light of the unified mechanistic framework proposed here, a major next step will be to determine how this shared CYTOR/MORRBID program contributes to normal cell-state transitions, tissue remodeling, and nonmalignant disease, and to what extent regulation of ERK signaling represents a common organizing principle across these settings.

This broader biological view also sharpens the therapeutic question. CYTOR has already been advanced as a biomarker and potential target in cancer ^34^, and proof-of-principle studies have shown both that CYTOR-directed RNA targeting can suppress metastatic behavior in vivo ^66^ and that delivery of a functional CYTOR RNA domain can improve dystrophic myotube function ^63^.

However, our findings argue that effective therapeutic development will require more than suppressing transcript abundance alone. It will require molecular resolution of the CYTOR/MORRBID-protein interaction itself: whether MEK2 binding is direct, which RNA elements and isoforms are required, how locus-specific transcripts modify this interaction in particular cellular contexts, and how this complex influences kinase activation or signaling-complex assembly. Resolving these questions will be essential for defining the therapeutic window of targeting this axis and for determining whether the shared CYTOR/MORRBID program represents a selective vulnerability in cancer or a broader regulator of tissue biology.

## Conclusions

Taken together, CYTOR and MORRBID provide a powerful model for studying lncRNA biology across evolutionary, mechanistic, and translational scales. The diverse and sometimes conflicting functions assigned to these loci should not be mistaken for an exhaustive account of their biology, nor should they discourage deeper mechanistic investigation. Instead, they underscore that lncRNA function is layered, context-dependent, and still incompletely resolved. In this regard, the CYTOR/MORRBID pair provides a tractable framework for defining how duplicated lncRNAs preserve a core functional output, how additional isoforms may contribute under specific cellular conditions, and how cytoplasmic lncRNA-protein interactions can be resolved at molecular detail. More broadly, this system illustrates why mechanistic understanding of lncRNAs remains an open and experimentally accessible problem with implications for both basic biology and therapeutic development.

## Methods

### Cell culture

Biological Industries MDA-MB-231, U2OS, MCF7, and HEK293 cells were maintained in Dulbecco’s Modified Eagle Medium (DMEM) supplemented with 10% fetal calf serum (FCS), 1% L-glutamine, and 1% penicillin–streptomycin. H1299 cells were cultured in RPMI 1640 medium (Gibco) supplemented with 10% FCS, 1% L-glutamine, and 1% penicillin–streptomycin (Biological Industries

### sgRNA design

Single-guide RNAs (sgRNAs) were designed using the CHOPCHOP web tool. Two sgRNAs targeting the lncRNA CYTOR were selected: sgRNA1 (GAGGGAAATAAATGACTGGA) and sgRNA3 (GTGTACATCATTGGGAATGG). A non-targeting sgRNA, negative control (NC) (GTCCACCCTTATCTAGGCTA), which does not target any sequence in the human genome, was used as a negative control.

### Cloning of sgRNA sequences into plasmids

Protospacer sequences were cloned into the sgRNA backbone plasmid pSB700, a gift from George Church (Addgene plasmid #64046), which was available with either puromycin or blasticidin selection markers. Approximately 100 ng of plasmid DNA was digested with the appropriate restriction enzymes: Esp3I for pSB700 plasmids and Bpu1102I and BstXI for pCRISPRia-v2 constructs. Annealed sgRNA oligonucleotides were ligated into the digested plasmids using T4 DNA ligase (New England Biolabs).

Ligation products were transformed into competent E. coli DH5α cells by heat shock and selected on ampicillin-containing plates. Individual colonies were expanded, and plasmid DNA was isolated using the Plasmid DNA Miniprep Kit (Bio Basic) according to the manufacturer’s instructions. Correct insertion of sgRNA sequences was confirmed by Sanger sequencing performed by Macrogen.

### Lentiviral transfection and transduction of CRISPRi system and sgRNAs

Lentiviral transfection was performed using Polyjet In Vitro Transfection Reagent (SignaGen, Frederick, MD, USA) following the manufacturer protocol. HEK293 cells were seeded 24 h before transfection to achieve approximately 90% confluency at the time of transfection. Cells were co-transfected with the plasmid of interest together with the lentiviral packaging plasmids VSVG (Addgene plasmid #8454) and psPAX2 (Addgene plasmid #12260).

48 hours after transfection, viral supernatants were collected and used for lentiviral transduction in the presence of Polybrene Infection Reagent (Sigma-Aldrich). Naive cells were first transduced with lentiviral particles carrying the dCas9-KRAB-mCherry construct. mCherry-positive cells were isolated using the BD FACSAria IIIU Cell Sorter and the Bigfoot Cell Sorter. Cells stably expressing dCas9-KRAB-mCherry were subsequently transduced with lentiviral particles carrying individual sgRNAs, with one sgRNA used per reaction. GFP-expressing plasmids were included as positive controls for transfection and transduction efficiency, and GFP expression was confirmed by fluorescence microscopy. Stable transductants were selected using blasticidin or puromycin, depending on the resistance marker encoded by the vector

### RNA-seq library prep and sequencing

#### 1. RNA QC

Quality and concentration measurements for total RNA were performed using the TapeStation 4200 (Agilent) with the RNA kit (Agilent, cat no. 5067-5576). All samples exhibited high integrity (RIN 10, excluding sample ’ACE A1’ with RIN 7.7).

#### 2. Library preparation

26 RNA-seq libraries were constructed simultaneously using NEBNext UltraExpress RNA Library Prep Kit for Illumina (NEB, cat no. E3330). 100 ng total RNA was used as starting material. mRNAs pull-down was performed using the NEBNext® Poly(A) mRNA Magnetic Isolation Module (NEB, cat no. E7490). RNA-seq library QC was performed by measuring library concentration using Qubit (Invitrogen) with the Equalbit dsDNA HS Assay Kit (Vazyme, cat no. EQ121) and size determination using the TapeStation 4200 (Agilent) with the High Sensitivity D1000 kit (Agilent, cat no. 5067-5584).

#### 3. Sequencing Details

The RNA-seq data was generated on Illumina NextSeq2000, using P4 XLEAP-SBS Reagent Kit (50 cycles) (Read1-72; Index1-8; Index2-8) (Illumina, cat no. 20100995).

### RNA-seq Analysis

RNA-seq data analysis was performed using the Galaxy platform. Raw sequencing reads in FASTQ format were uploaded directly to Galaxy and assessed for quality using FastQC. Adapter sequences and low-quality bases were removed using Trimmomatic. Cleaned reads were aligned to the human reference genome GRCh38 using HISAT2 with strand-specific parameters. Gene annotation files in GTF format were obtained from GENCODE. Aligned reads were quantified at the gene level using featureCounts based on the corresponding genome annotation.

Downstream analyses were performed using count matrices generated from uniquely mapped reads. Followed by downstream analyses in R. Differential expression analysis between the samples and negative controls was performed using the DESeq2 package. Differentially expressed genes were defined as those with a log₂ (fold change) greater than 1 or less than -1, with an adjusted p-value of less than 0.05.

### Apoptosis

A day before the assay, cells were seeded at a density of 1 x 106 cells per reaction in 2mL medium in a 6-well plate. Cells were washed twice by phosphate-buffer saline (PBS), and resuspended with 100uL of Annexin V Binding Buffer (BioLgened). Cells were incubated for 15 mins in the dark with 3ul of FITC Annexin V (Biolegend), 2ug of DAPI. Samples were then diluted with 200uL Annexin V Binding Buffer. Cells were analyzed by the Aurora Spectral Flow Analyzer

### Cell cycle

For cell cycle analysis, 6–7 × 10^5 HCT116 cells were seeded per well in 6-well plates, with three biological replicates per condition. Cells were washed once with PBS and fixed in 70% cold ethanol for at least 30 min at 4 °C. Fixed cells were centrifuged at 2,000 rpm for 5 min, washed once with PBS, and stained with DAPI at a final concentration of 1 μg/mL for 5 min. Samples were analyzed by flow cytometry NovoCyte analyzer.

### Growth assay and Crystal violet

For growth assays, 5,000 cells per well were seeded in 6-well plates in the appropriate culture medium. Cells were maintained in DMEM or RPMI depending on the cell line used. At each time point, one plate was collected for crystal violet staining while the remaining plates received fresh medium. For crystal violet staining, culture medium was removed and cells were washed once with ice-cold PBS. Cells were fixed with 1 mL ice-cold methanol per well for 20 min at room temperature. Methanol was removed and cells were stained with 1 mL of 0.25% crystal violet solution for at least 20 min at room temperature with gentle shaking. Wells were washed thoroughly with water to remove excess stain, and plates were inverted on filter paper and air-dried without lids for at least 2 h at room temperature. For optimal drying, plates were left at room temperature for 16–24 h. For quantification, 1 mL of 30% acetic acid was added to each well and incubated for 20 min at room temperature with gentle shaking to solubilize the bound dye. Absorbance was measured at 595 nm, and blank values were subtracted from cell-containing wells. OD595 values were used as a quantitative measure of relative cell growth.

### Protein extraction and western blotting

Proteins were extracted using RIPA lysis buffer according to the manufacturer’s instructions. Protein concentrations were determined using the BCA assay. Equal amounts of protein were used for western blot analysis. Membranes were incubated with the following primary antibodies: MEK2 (ab178876), phospho-MEK1/2 (#9121), p44/42 MAPK (ERK1/2; #9102), phospho-p44/42 MAPK (ERK1/2; #9101), and GAPDH (#2118). Membranes were subsequently incubated with goat anti-rabbit IgG secondary antibody (#7074). All antibodies were obtained from Cell Signaling Technology, except MEK2 (abcam).

### RNA extraction

Total RNA was extracted using TRI Reagent according to the manufacturer’s instructions. Briefly, cell pellets were lysed in 1 mL TRI Reagent and incubated for 5 min at room temperature. Chloroform (200 μL) was added to each sample, followed by vigorous mixing and centrifugation at >12,000 × g for 15 min at 4 °C to separate the aqueous and organic phases. The upper aqueous phase was transferred to a fresh tube, mixed with 1 μL glycogen blue and 500 μL isopropanol, and incubated for 5–10 min at room temperature to precipitate RNA. Samples were centrifuged at >12,000 × g for 10 min at 4 °C, and RNA pellets were washed with 75% ethanol, air-dried, and resuspended in nuclease-free water. Purified RNA was either used immediately for cDNA synthesis or stored at −80 °C

### Quantitative PCR (qPCR)

Quantitative PCR (qPCR) was performed using qPCRBIO SyGreen Master Mix (PB20.15-50) on a QuantStudio real-time PCR system according to the manufacturers’ instructions.

### Reverse Transcription (RT)

Complementary DNA (cDNA) was synthesized from purified RNA using the qScript cDNA Synthesis Kit according to the manufacturer’s instructions.

### Cross-linking immunoprecipitation followed by qPCR (CLIP–qPCR)

Cells (20–30 × 10^6) were washed with ice-cold PBS and irradiated with UV light at 254 nm to crosslink RNA-protein complexes. For MDA-MB-231 cells, a UV dose of 150 mJ/cm^2 was used. Following irradiation, cells were scraped, collected by centrifugation, and either processed immediately or snap-frozen in liquid nitrogen and stored at −80 °C. Cell pellets were lysed in IP lysis buffer containing 20 mM HEPES (pH 7.4), 150 mM NaCl, 10% glycerol, 1% Triton X-100, and 1 mM EGTA supplemented with protease and RNase inhibitors. Lysates were incubated on ice for 30 min, clarified by centrifugation at 15,000 rpm for 10 min at 4 °C, and protein concentrations were determined. For each immunoprecipitation, approximately 1 mg of total protein was used. A fraction of the lysate was retained as an input control for RNA and protein analysis. Lysates were incubated with 3 μg MEK2 antibody (A-1; sc-13159) for 4 h at 4 °C, followed by overnight incubation at 4 °C with Protein G magnetic beads (Dy-10003D; Thermo Fisher Scientific). No-antibody and IgG isotype controls were included in parallel. Beads were washed three times with IP lysis buffer and treated with TURBO DNase (AB-AM2238; Thermo Fisher Scientific) at 37 °C for 30 min. After additional washes, 10% of each immunoprecipitated sample was reserved for protein analysis, and the remaining material was incubated with proteinase K at 55 °C for 30–60 min. RNA was extracted using TRIzol reagent followed by chloroform separation and purification with the RNA Clean & Concentrator-5 kit. Purified RNA was eluted in nuclease-free water and stored at −80 °C until use. Reverse transcription was performed using random hexamer primers and SuperScript III reverse transcriptase (18080044; Thermo Fisher Scientific). Complementary DNA was diluted and analyzed by quantitative PCR.

### EPHA4 Overexpression

EPHA4 overexpression constructs were generated by cloning the coding sequence of EPHA4 (ENST00000281821.7; 2,969 bp) into the N174 #81061 expression vector. The EPHA4 insert was synthesized by Twist, and added to the both side primer sequence. Both the insert and N174 plasmid were digested with NotI and MluI restriction enzymes according to the manufacturer’s instructions. Digested products were validated by agarose gel electrophoresis and purified prior to ligation. Insert and vector fragments were ligated using T4 DNA ligase, and ligation reactions were transformed into competent bacteria. Positive colonies were screened by colony PCR and confirmed by Sanger sequencing performed by Macrogen.

### Single-molecule RNA fluorescence in situ hybridization (RNA FISH)

Single-molecule RNA fluorescence in situ hybridization (smiFISH) was performed on cells grown on coverslips in 24-well plates for at least 24 h before fixation according to this protocol ^67^. Briefly, Cells were washed with PBS and fixed with freshly prepared 4% paraformaldehyde for 10–20 min at room temperature, followed by quenching and permeabilization with 0.1% Triton X-100 in RNase-free PBS for 10 min. Samples were then incubated in pre-hybridization buffer containing formamide for 15 min at room temperature. Probe–FLAP hybridization mixtures were prepared in hybridization buffer and applied to parafilm droplets in humidified chambers. Coverslips were inverted onto the hybridization mixture and incubated overnight at 37 °C protected from light. The following day, coverslips were washed twice with warm formamide buffer at 37 °C for 30 min, followed by washes in 1× SSC buffer and PBS. Samples were mounted, sealed with nail polish, and stored at 4 °C protected from light until imaging.

### RNA-sequencing analysis of publicly available datasets for MORRBID/CYTOR expression

Normalized ene expression data (STAR-aligned, log2(TPM+1)-transformed) and corresponding clinical annotations were obtained from the UCSC Xena platform for TCGA and TARGET cohorts. Ensembl gene identifiers were mapped to HGNC gene symbols using the org.Hs.eg.db annotation package (v3.18). Duplicate gene symbols were removed, retaining the first occurrence. MORRBID and CYTOR expression were compared between primary tumor samples (TCGA barcode -01) and adjacent solid tissue normal samples (-11) using the Wilcoxon rank-sum test. For TCGA-BRCA, analysis was restricted to female patients. MORRBID and CYTOR expression were compared by using Wilcoxon signed-rank test.

### Gene set enrichment analysis

Pearson correlation coefficients were computed between MORRBID and CYTOR expression and all other genes across tumor samples. The resulting ranked gene list was used as input for pre-ranked GSEA against the MSigDB Hallmark gene set collection, using the clusterProfiler R package. Pathways with Benjamini-Hochberg adjusted p-values < 0.05 were considered significant. GSEA was performed separately in tumor and normal tissue samples.

### Survival Analysis

Kaplan-Meier overall survival analysis was performed on tumor samples. The optimal expression cutoff for dichotomizing patients into high and low expression groups was determined using the maximally selected rank statistic via the surv_cutpoint() function (survminer R package). Survival times were converted from days to months. Survival curves were compared using the log-rank test, with follow-up truncated at 60 months.

## Abbreviation

lncRNAs: Long non-coding RNAs
RNA-seq: RNA sequencing
GTEx: Genotype-Tissue Expression
CRISPRi: CRISPR interference
PCGs: Protein-coding genes
EMT: Epithelial–mesenchymal transition
RT-qPCR: Reverse transcription–quantitative PCR
EPHA4: Ephrin type-A receptor 4
RAP-MS: RNA antisense purification–mass spectrometry
MEK2 (MAP2K2): Mitogen-activated protein kinase kinase 2
smRNA-FISH: Single-molecule RNA fluorescence in situ hybridization
CLIP-qPCR: Cross-linking immunoprecipitation followed by qPCR
TRE: TPA response element
SRE: Serum response element NC Negative control

## Author Contributions

T.A.N., S.A.M., and N.L.J. conducted experiments and performed analyses. O.B., Z.Y., and D.A. performed computational analyses. T.A.N. and A.C.B. wrote the manuscript. A.C.B. conceived and supervised the study.

## Acknowledgements

The authors thank Prof. Alejandro Chavez (University of California San Diego) for the pSB700 plasmids. The authors thank Prof. Gregor Neuert (Vanderbilt University) for helping with smRNA-FISH. The authors gratefully acknowledge the LS&E Microscopy and Flow Cytometry Units. The authors gratefully acknowledge the Azrieli Technion Genomics Center (ATGC).

