## Supplementary figures and images for "Paralogous lncRNAs CYTOR and MORRBID share a conserved trans acting function in MEK ERK signaling"

### Supplementary fig 2

Supplementary Figure.2

a

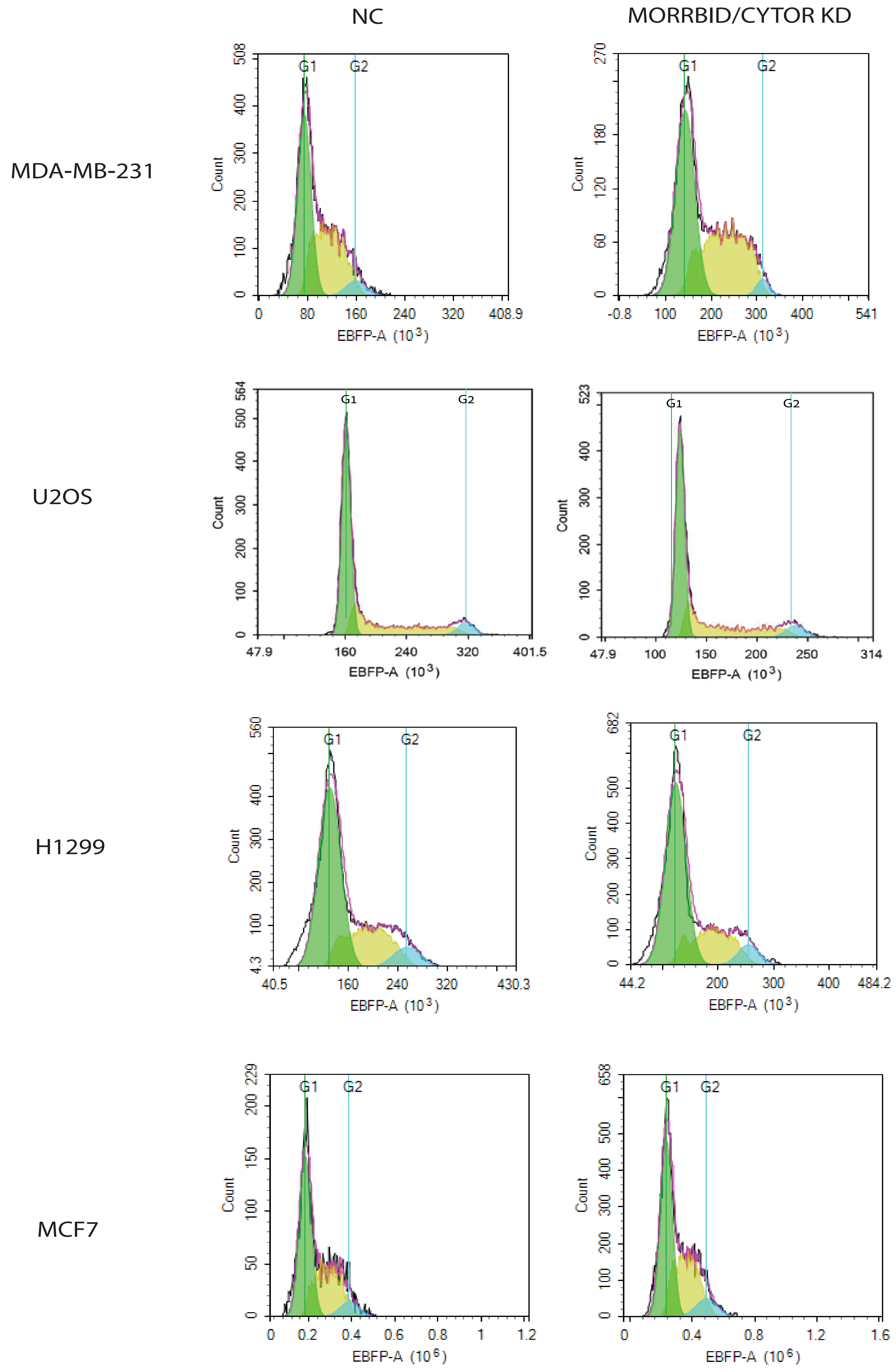

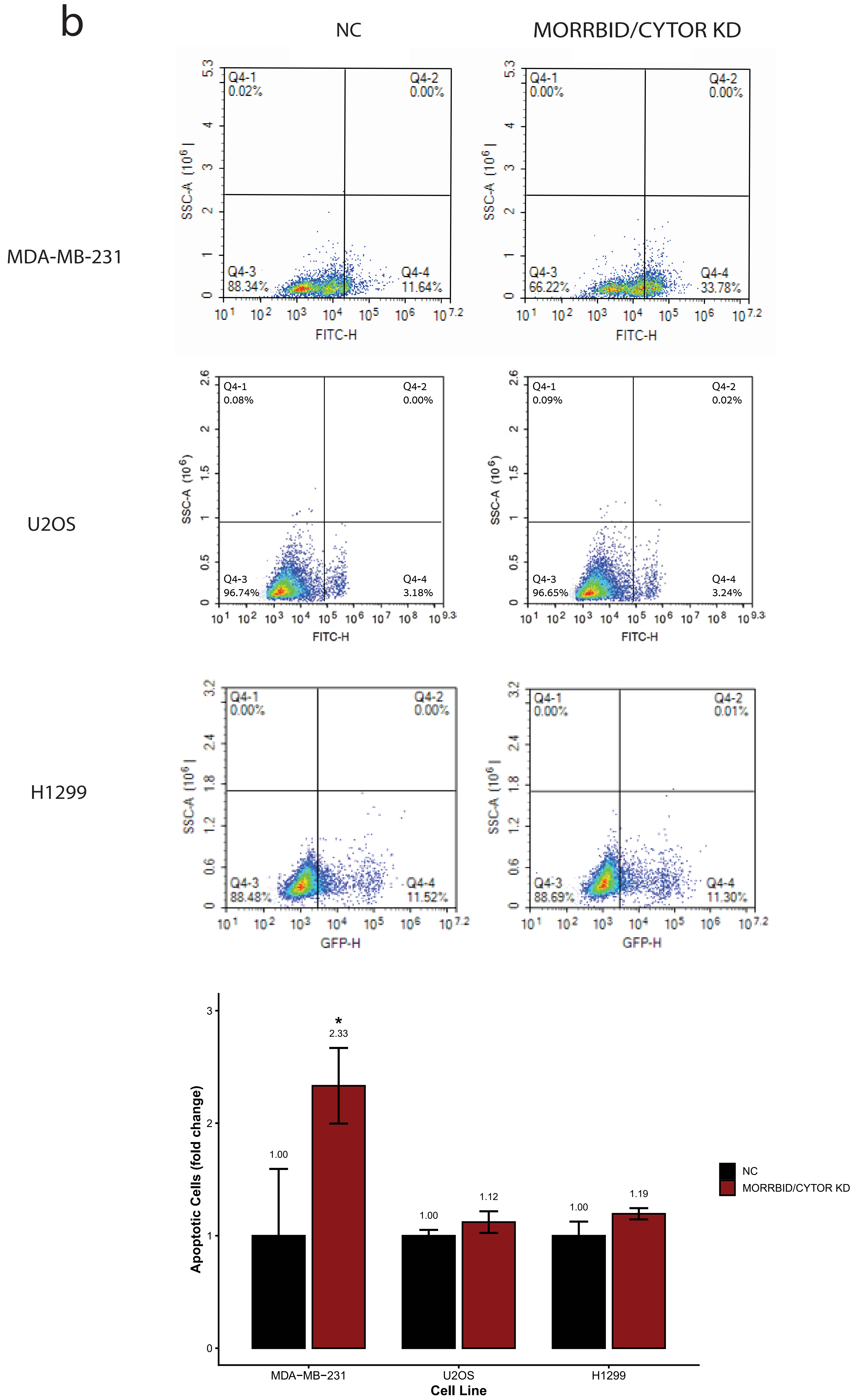

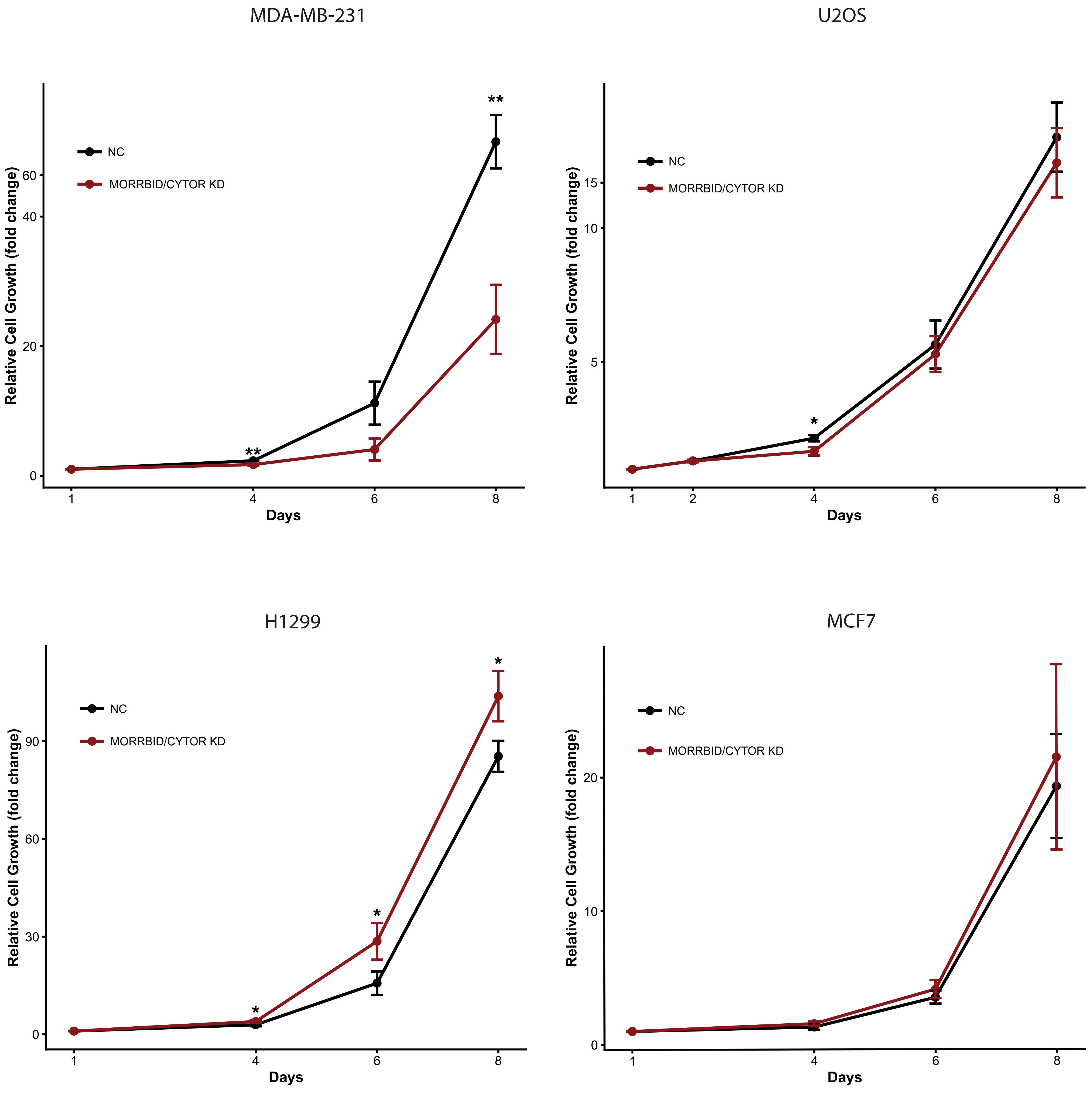

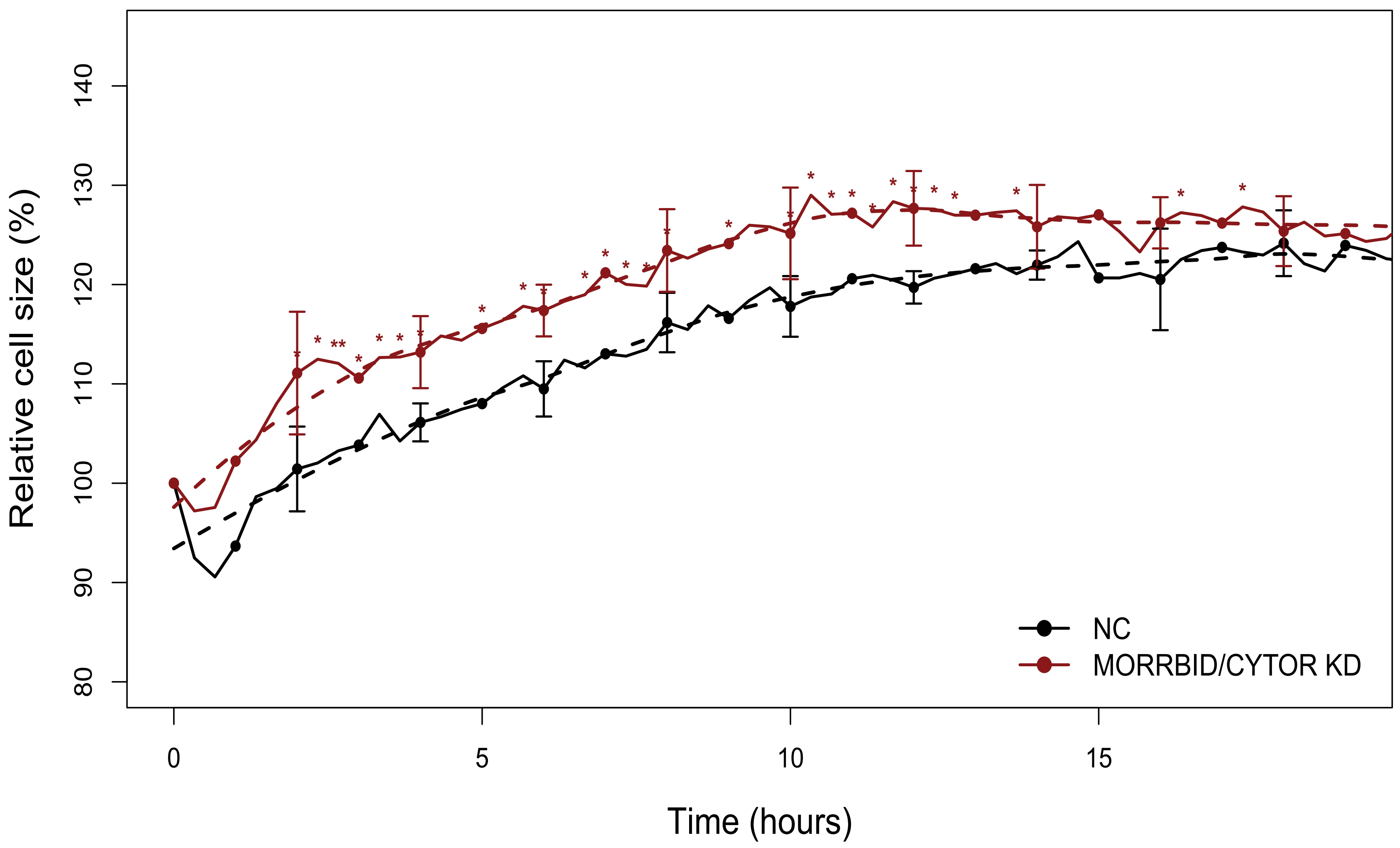

### Supplementary fig 3

Supplementary Figure.3

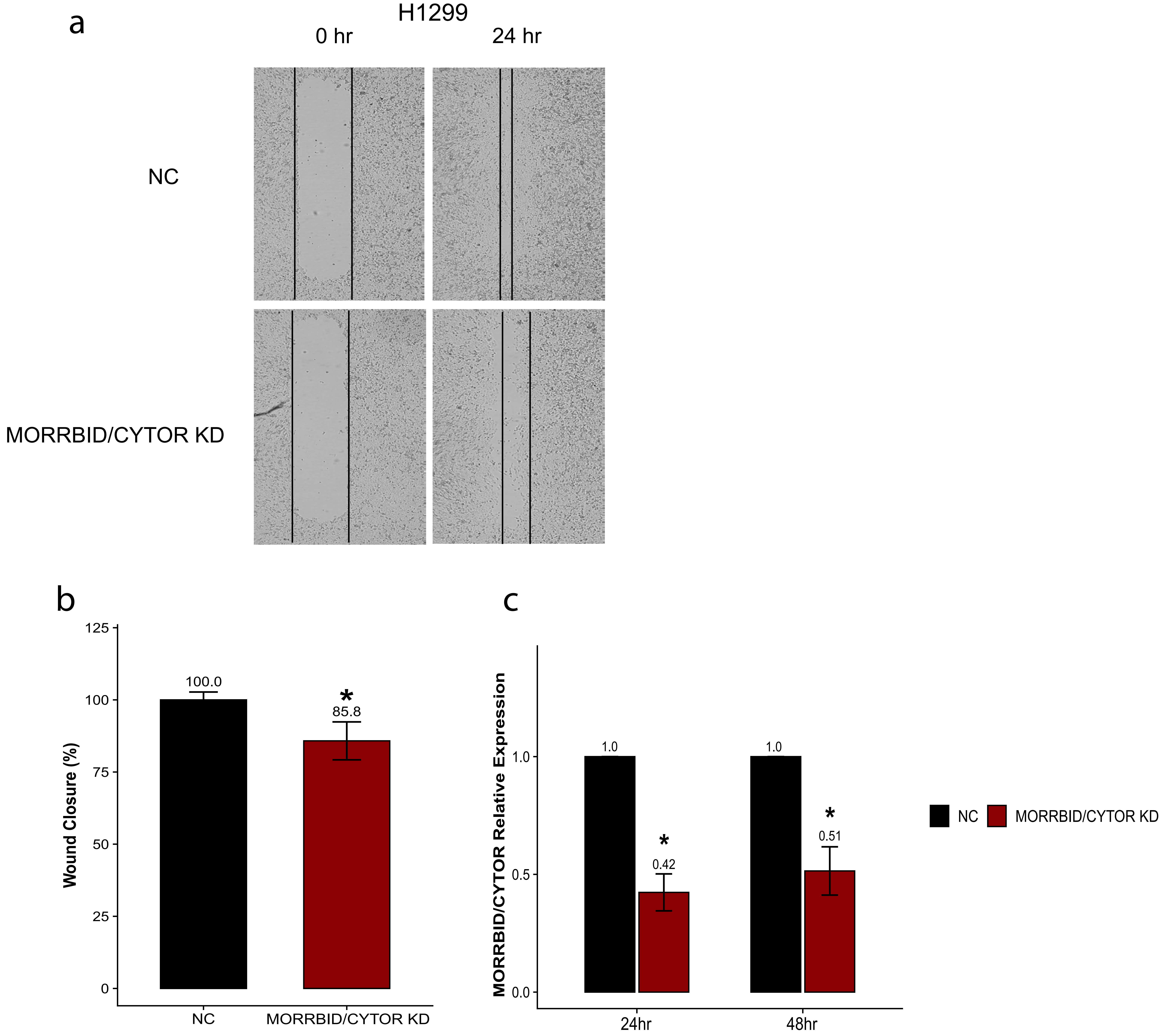

### Supplementary fig 4

Supplementary Figure.4

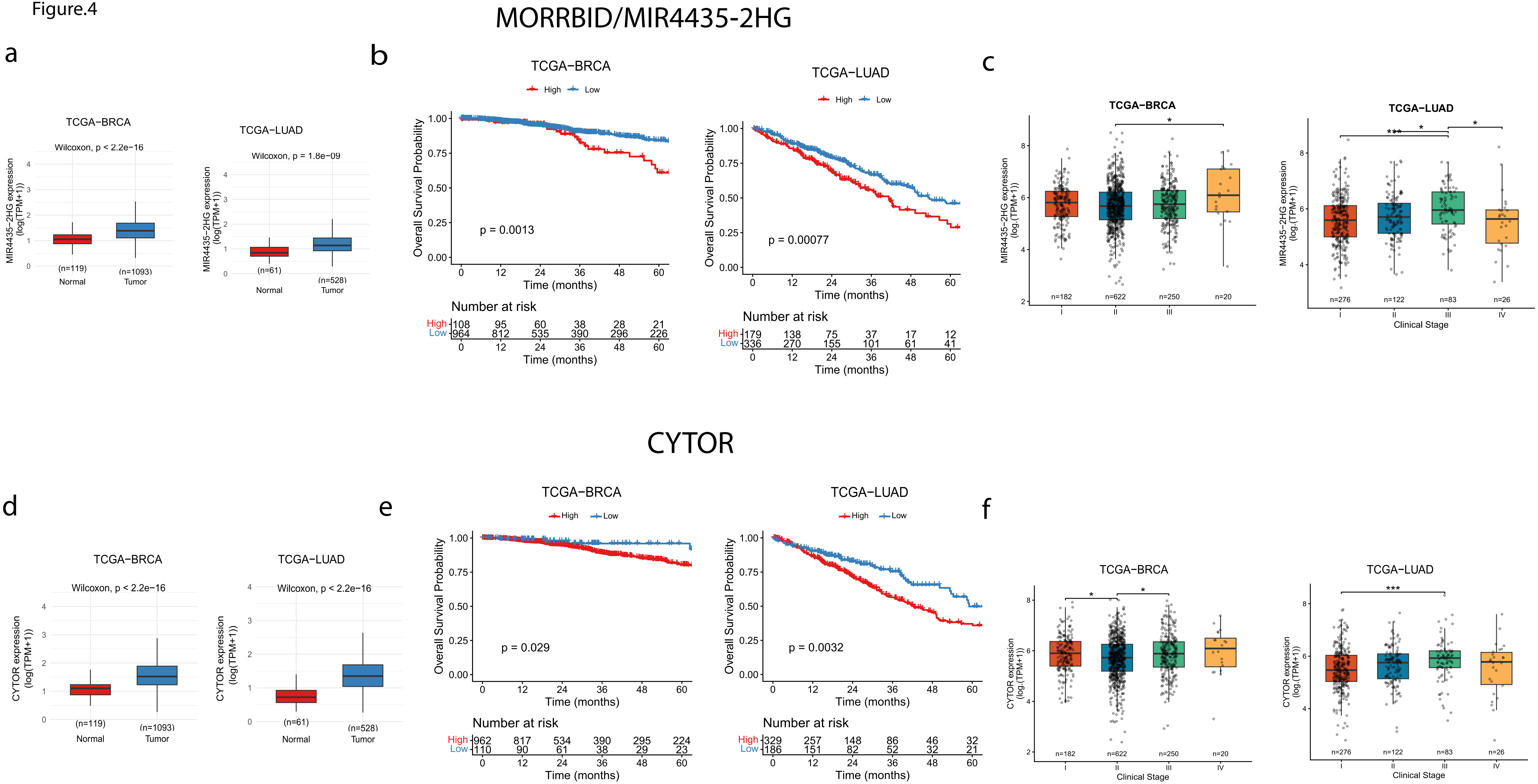

### Supplementary fig 5

Supplementary Figure.5

a

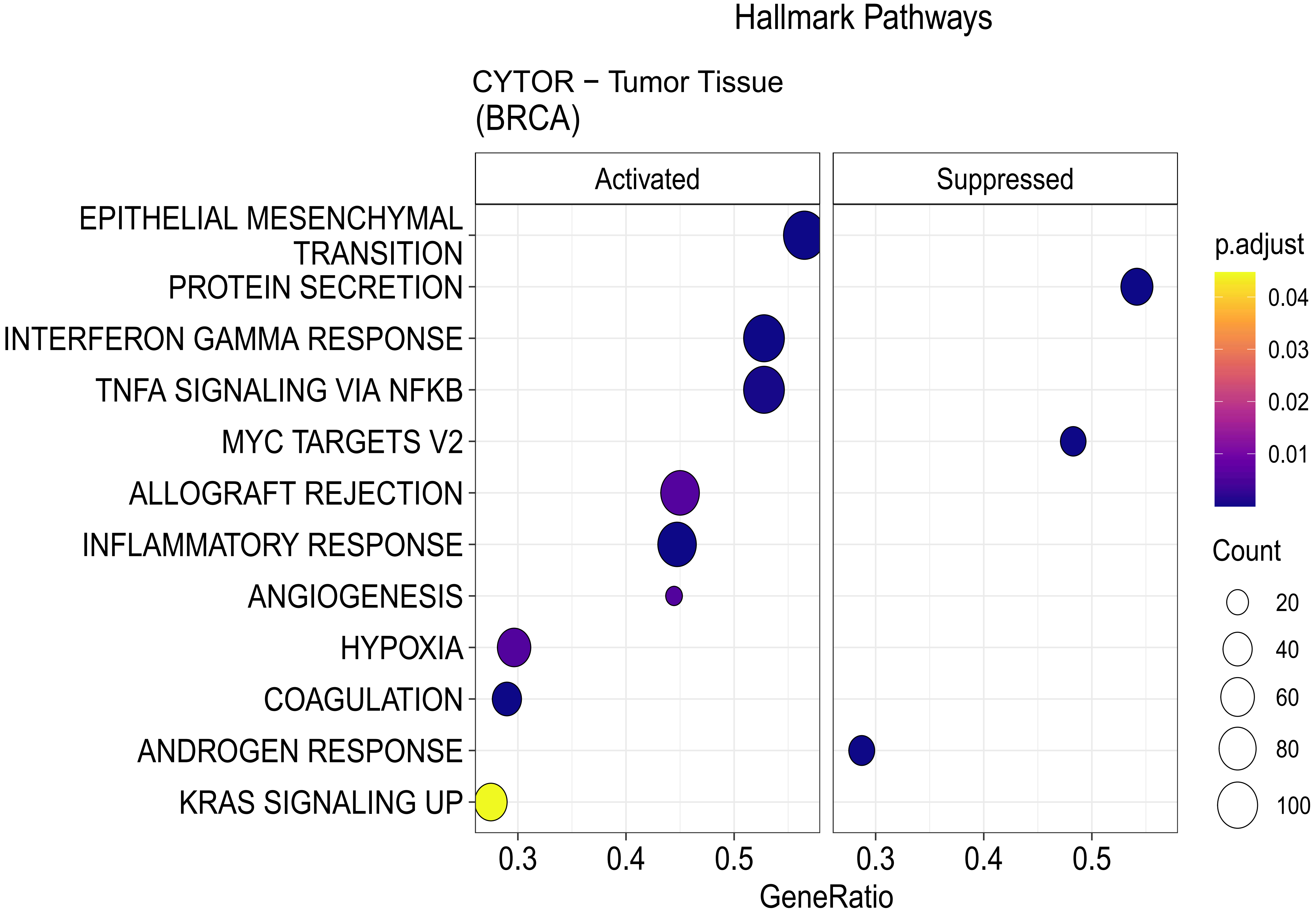


b

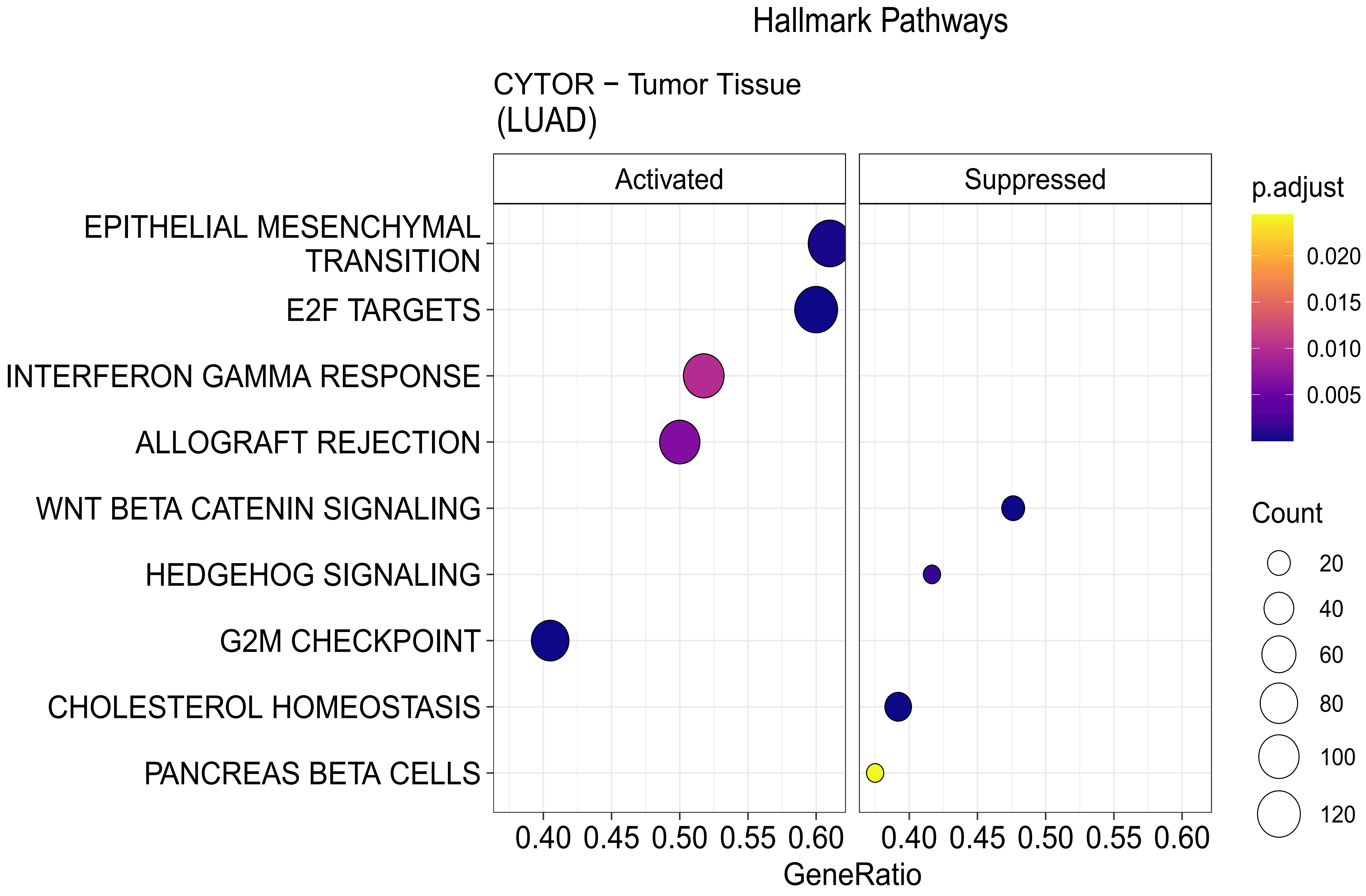

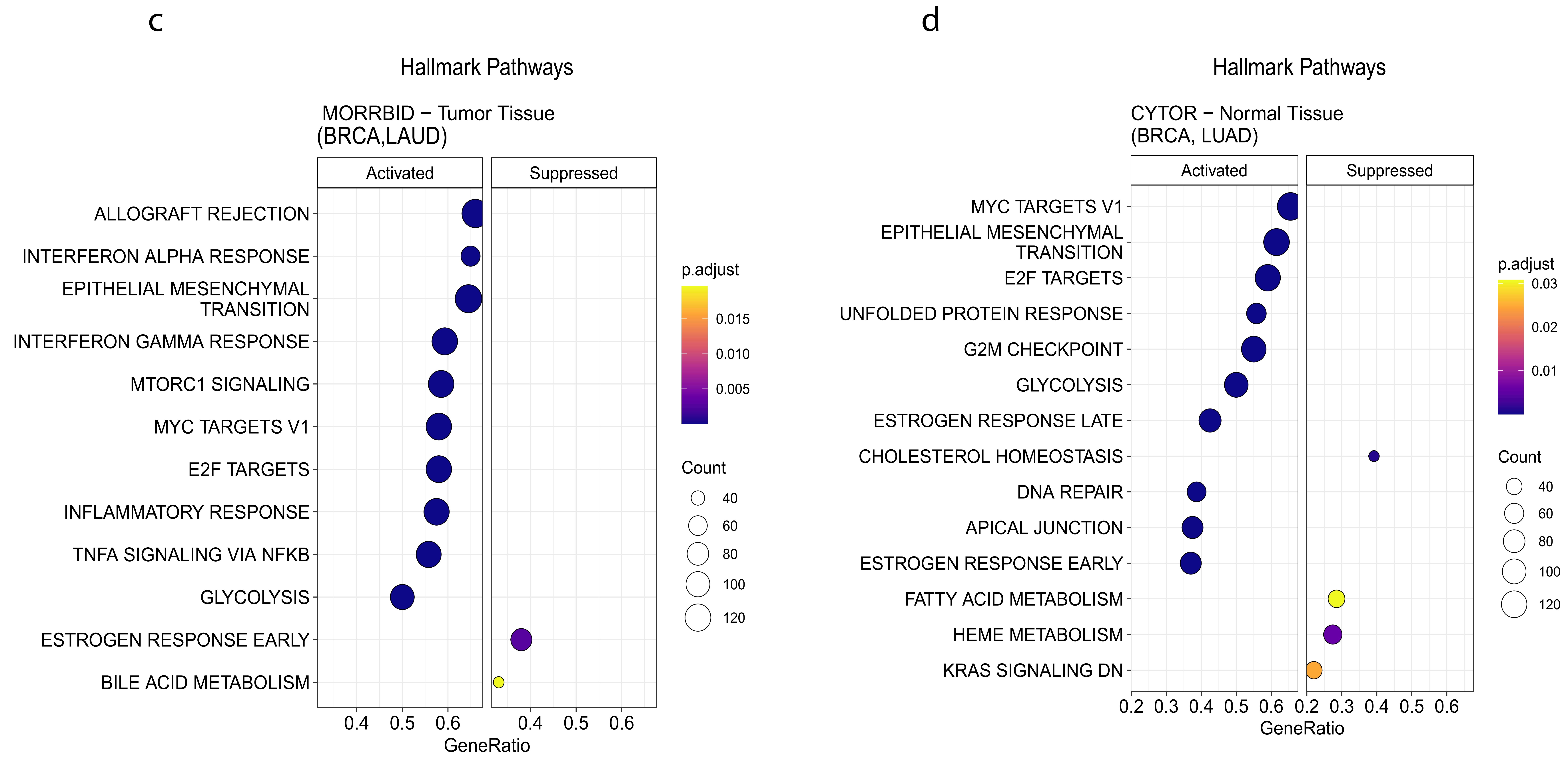

### Supplementary fig 6

Supplementary Figure.6

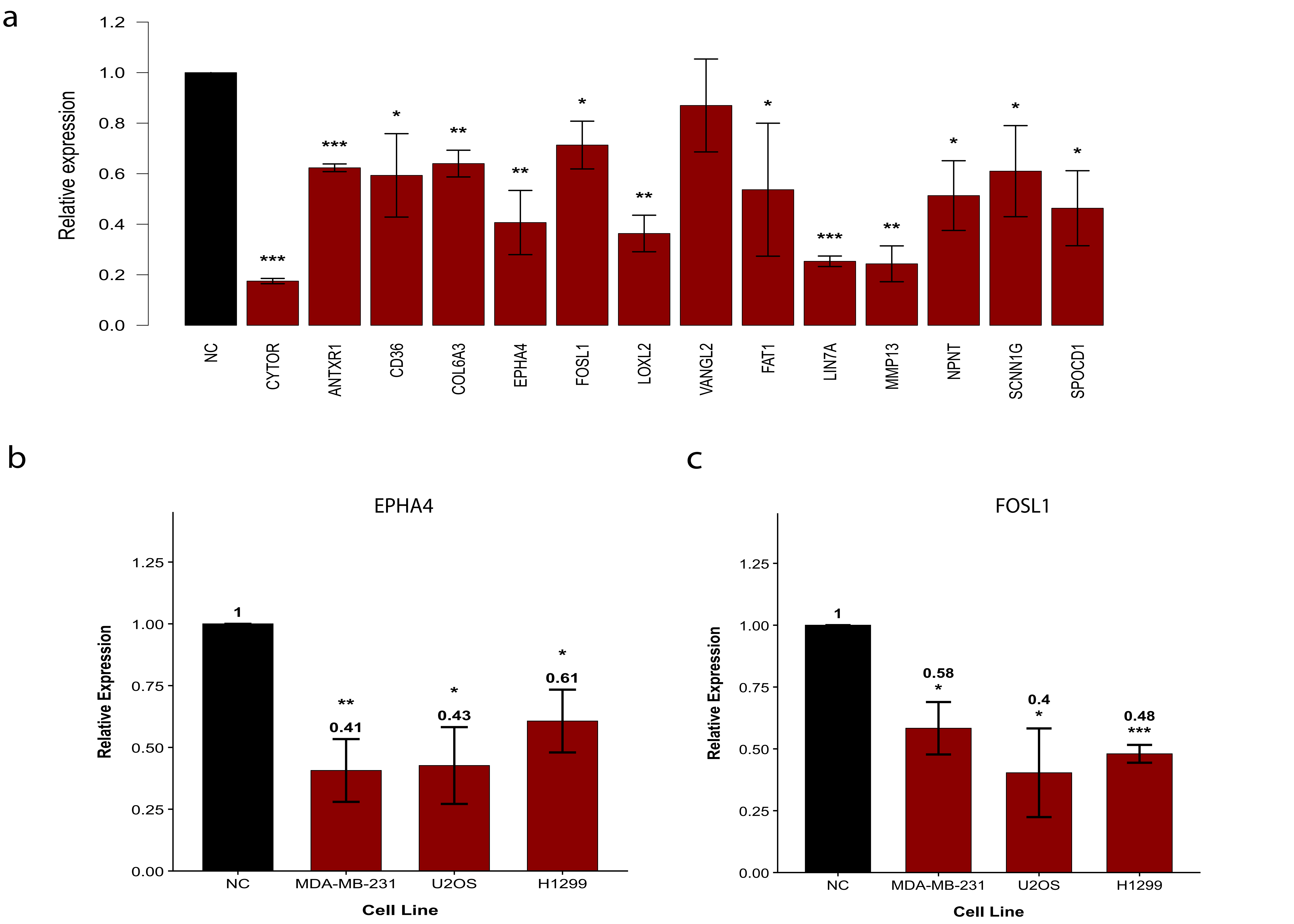

### Supplementary fig 7

Supplementary Figure.7

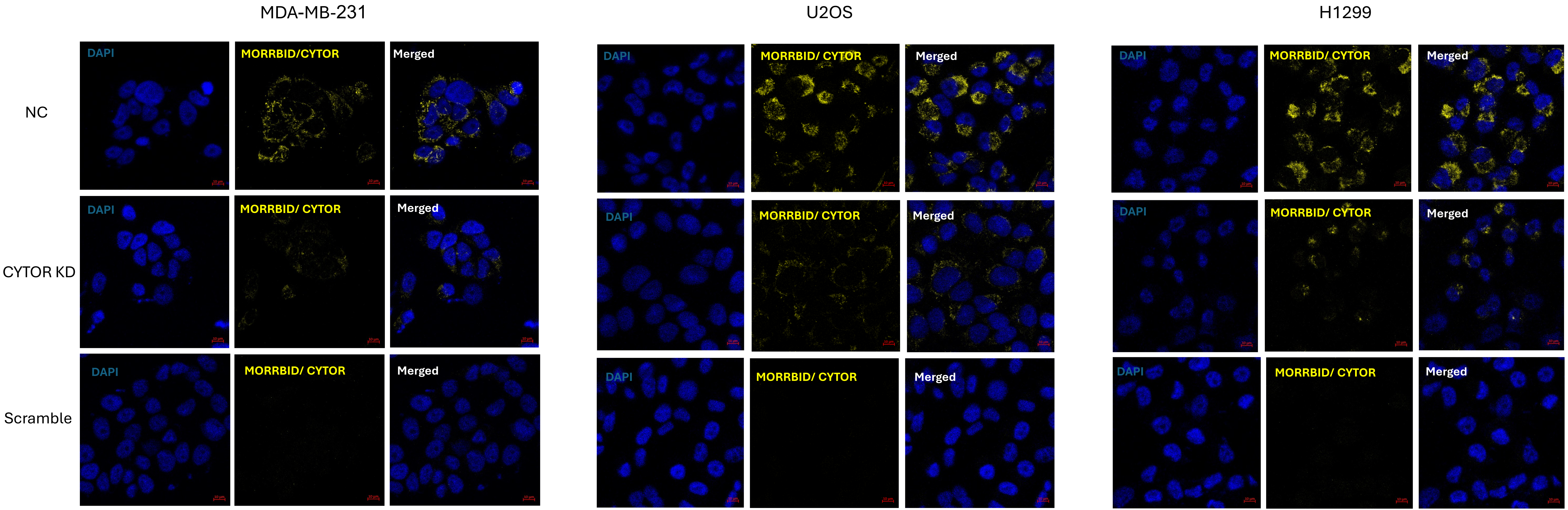

### Supplementary fig 8

Supplementary Figure.8

a

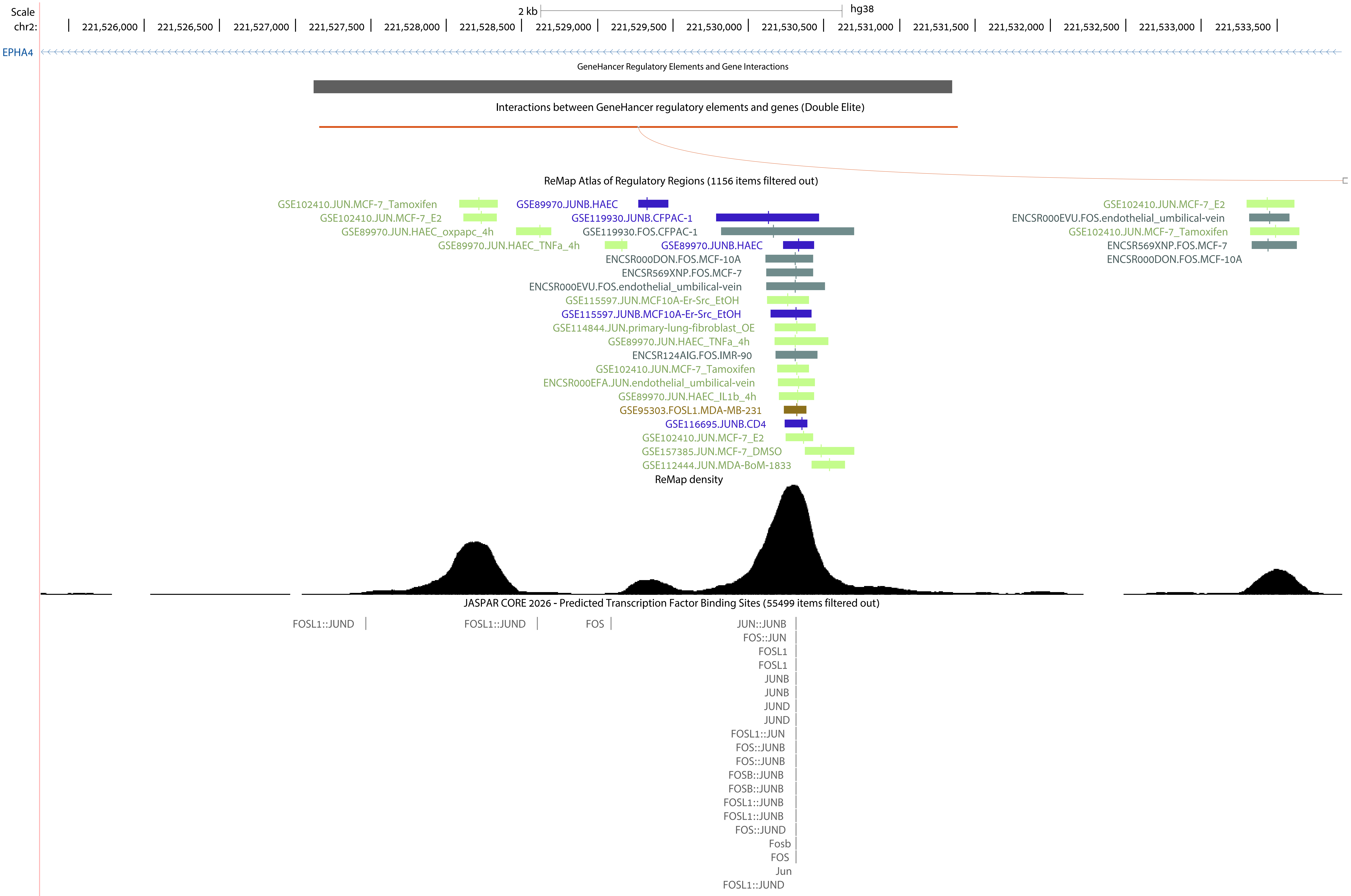

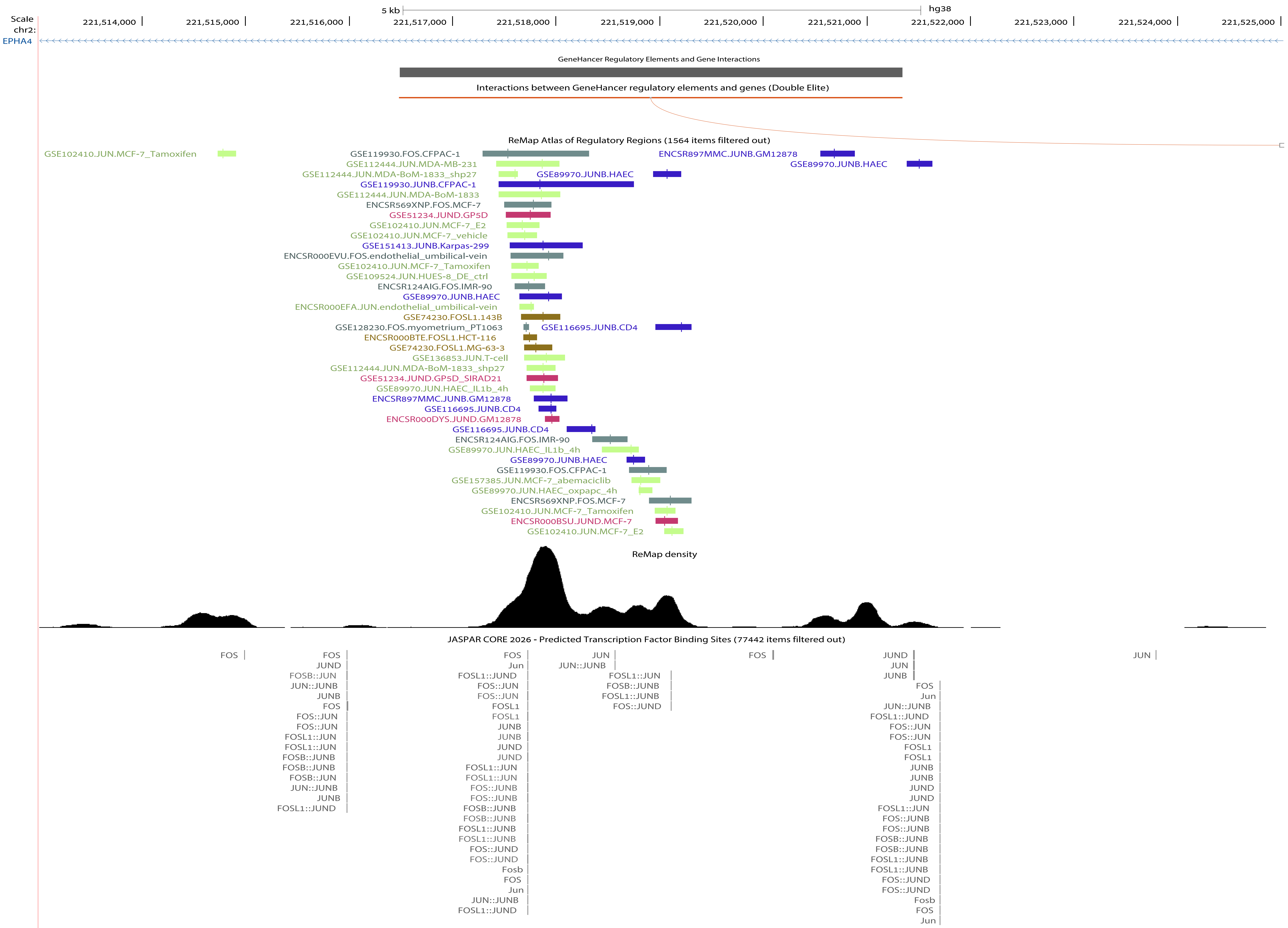

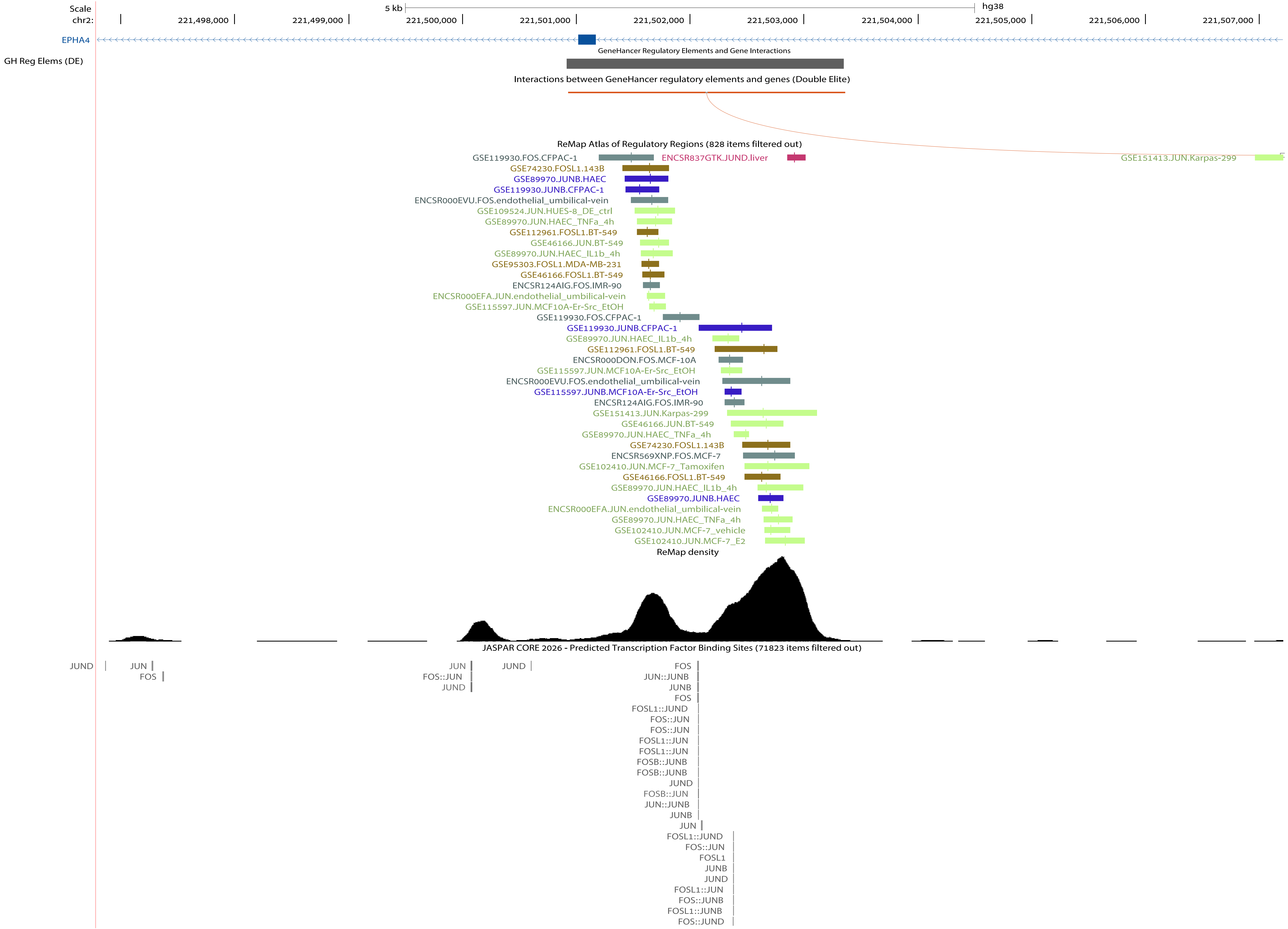
