## Supplementary figure legends for "Paralogous lncRNAs CYTOR and MORRBID share a conserved trans acting function in MEK ERK signaling"

**Supplementary Figure Legends**
 **Fig. S1 MORRBID is associated with mesenchymal cell identity, extracellular matrix organization, and cancer-related transcriptional programs.**
**a** Mean MORRBID expression across cell types from CELLxGENE single-cell RNA-sequencing (scRNA-seq) datasets, shown as Z-scores of mean expression across cell types. **b** Gene Ontology (GO) Biological Process enrichment analysis of genes co-expressed with MORRBID in GTEx bulk RNA-sequencing (RNAseq) data. **c** Correlation analysis of CYTOR and MORRBID expression across non-diseased GTEx tissues showing a strong positive association between the two transcripts. **d** MORRBID expression across tumor samples and paired normal tissues from TCGA datasets. MORRBID expression is elevated in multiple tumor types relative to matched normal tissues. **e** CYTOR expression across tumor samples and paired normal tissues from TCGA datasets. CYTOR expression is elevated across multiple tumor types relative to matched normal tissues. **f** Correlation analysis of CYTOR and MORRBID expression across TCGA tumor samples showing a strong positive association between the two transcripts in cancer.

**Fig. S2 Effects of simultaneous CYTOR and MORRBID repression on cell cycle, apoptosis, proliferation, and cell spreading.
a** Cell-cycle analysis following simultaneous MORRBID/CYTOR knockdown (KD) in MDA-MB-231, U2OS, H1299, and MCF7 cells. **b** Apoptosis analysis following simultaneous MORRBID/CYTOR KD in MDA-MB-231, U2OS, and H1299 cells. Bars represent mean ±SD. n=3, * p value<0.05, ** p value<0.01. **c** Relative cell growth following simultaneous MORRBID/CYTOR KD in MDA-MB-231, U2OS, H1299, and MCF7 cells. Data are presented as mean ± SD. *P < 0.05, **P < 0.01 by Student’s t-test. **d** Quantification of relative cell size (%) over time following simultaneous MORRBID/CYTOR KD in MDA-MB-231 cells, normalized to the initial time point (t = 0). Data are presented as mean ± SD. *P < 0.05, **P < 0.01 by Student’s t-test.
**Fig. S3 siRNA-mediated knockdown of CYTOR and MORRBID impairs cell migration.**
**a** Representative images of wound healing assays in H1299 cells transfected with negative control (NC) or siRNAs targeting MORRBID/CYTOR KD at 0 and 24 hours. three biological replicates. **b** Quantification of wound closure (%) at 24 hours shows a significant reduction in migration upon CYTOR/MORRBID knockdown compared to NC. Bars represent mean ±SD. n=3, * p value<0.05. **c** RT-qPCR analysis confirms efficient knockdown of CYTOR and MORRBID at 24 and 48 hours post–siRNA transfection. Bars represent mean ±SD. n=2, * p value<0.05.

**Fig. S4 MORRBID/CYTO expression is associated with poor clinical outcome and advanced tumor stage.**
**a** MORRBID expression in paired normal and tumor samples from Breast Cancer and Lung Adenocarcinoma cohorts. MORRBID expression is significantly increased in tumor samples compared with normal tissues. **b** Kaplan–Meier analysis of overall survival in TCGA-BRCA and TCGA-LUAD cohorts grouped according to MORRBID expression. High MORRBID expression is associated with reduced overall survival. **c** MORRBID expression across clinical stages in TCGA-BRCA and TCGA-LUAD cohorts. MORRBID expression is increased in advanced-stage tumors. **d** CYTOR expression in paired normal and tumor samples from TCGA-BRCA and TCGA-LUAD cohorts. CYTOR expression is significantly increased in tumor samples compared with normal tissues. **e** Kaplan–Meier analysis of overall survival in TCGA-BRCA and TCGA-LUAD cohorts grouped according to CYTOR expression. High CYTOR expression is associated with reduced overall survival. **f** CYTOR expression across clinical stages in TCGA-BRCA and TCGA-LUAD cohorts. CYTOR expression is increased in advanced-stage tumors.

**Fig. S5 MORRBID/CYTOR-associated transcriptional programs are enriched for epithelial–mesenchymal transition pathways.**
 **a** Hallmark pathway enrichment analysis of genes co-expressed with CYTOR in breast tumor tissue. **b** Hallmark pathway enrichment analysis of genes co-expressed with CYTOR in lung adenocarcinoma tumor tissue. **c** Hallmark pathway enrichment analysis of genes co-expressed with MORRBID in breast and lung tumor tissue. **d** Hallmark pathway enrichment analysis of genes co-expressed with CYTOR in non-diseased breast and lung tissues.

**Fig. S6 Validation of MORRBID/CYTOR-regulated migration-associated genes.**
 **a** Reverse transcription–quantitative polymerase chain reaction (RT–qPCR) validation of selected migration-associated genes following simultaneous MORRBID/CYTOR KD in MDA-MB-231 cells. **b** RT-qPCR analysis of EPHA4 expression following simultaneous MORRBID/CYTOR KD in MDA-MB-231, U2OS, and H1299 cells. **c** RT-qPCR analysis of FOSL1 expression following simultaneous MORRBID/CYTOR KD in MDA-MB-231, U2OS, and H1299 cells. Data are presented as mean ± SD. *P < 0.05, **P < 0.01, ***P < 0.001 by Student’s t-test.

**Fig. S7 MORRBID/CYTOR is predominantly localized in the cytoplasm.**
 Representative smRNA FISH images showing MORRBID/CYTOR localization in MDA-MB-231, U2OS, and H1299 cells under negative control, MORRBID/CYTOR KD, and scramble probe conditions. DAPI marks nuclei.

**Fig. S8 AP-1 transcription factor occupancy at additional EPHA4 regulatory elements.**
 **a–c**, Zoomed views of the three additional EPHA4 intragenic enhancer regions identified by GeneHancer. ReMap ChIP–seq tracks show binding of JUN-, JUNB-, JUND-, FOS-, and FOSL1-family transcription factors across these enhancer elements, while JASPAR motif analysis identifies multiple predicted AP-1-related binding motifs, including FOSL1::JUN, FOSL1::JUND, FOS::JUN, FOS::JUNB, and FOS::JUND complexes.
