## Supplementary fig 1 for "Paralogous lncRNAs CYTOR and MORRBID share a conserved trans acting function in MEK ERK signaling"

Supplementary Figure.1

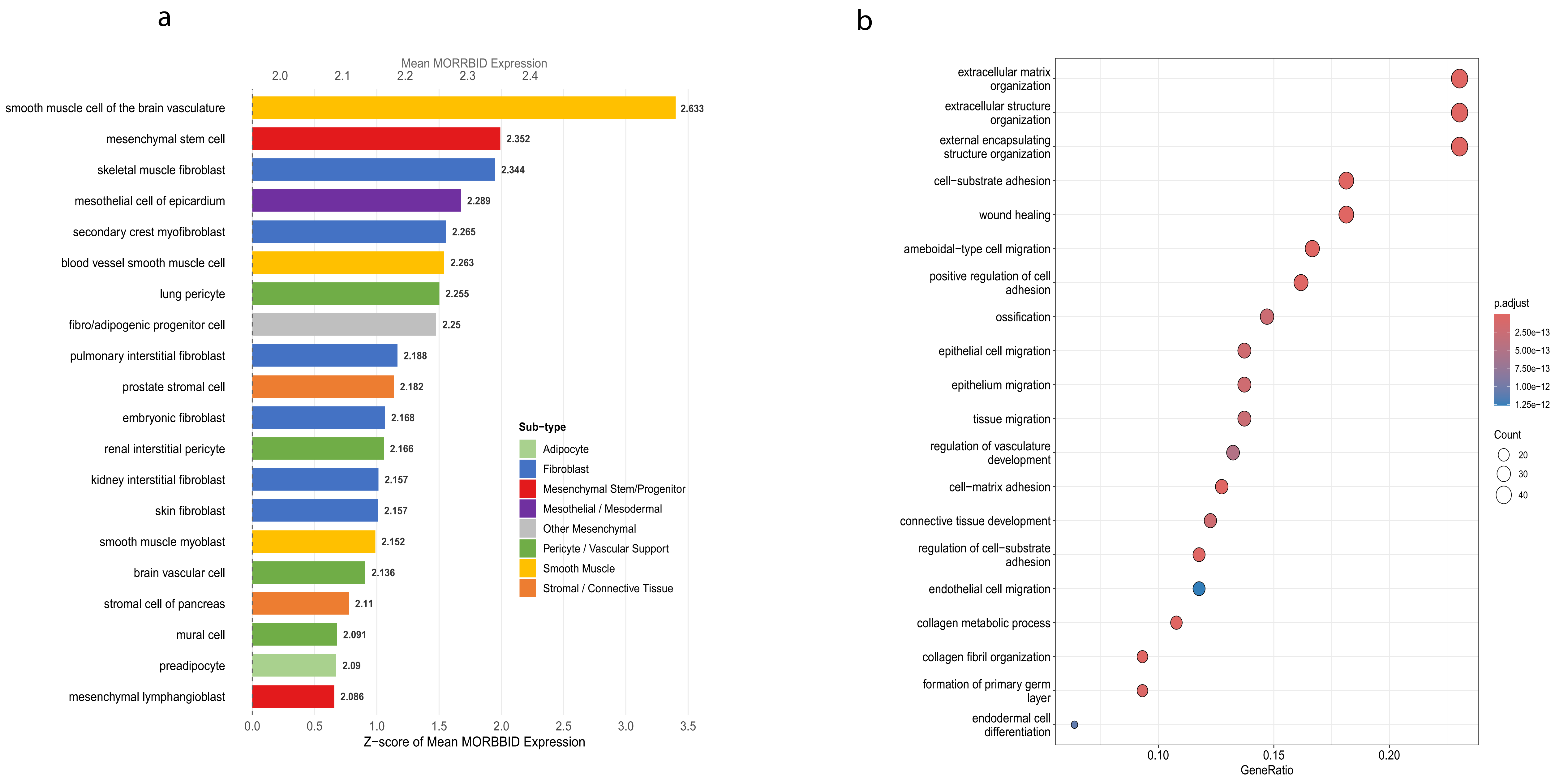

c

Correlation analysis result

| name | name | r | p-value | n_tumor | n_peritumor | n_gtex |
|---|---|---|---|---|---|---|
| CYTOR | MIR4435-2HG | 0.907 | 0 | 0 | 0 | 7862 |

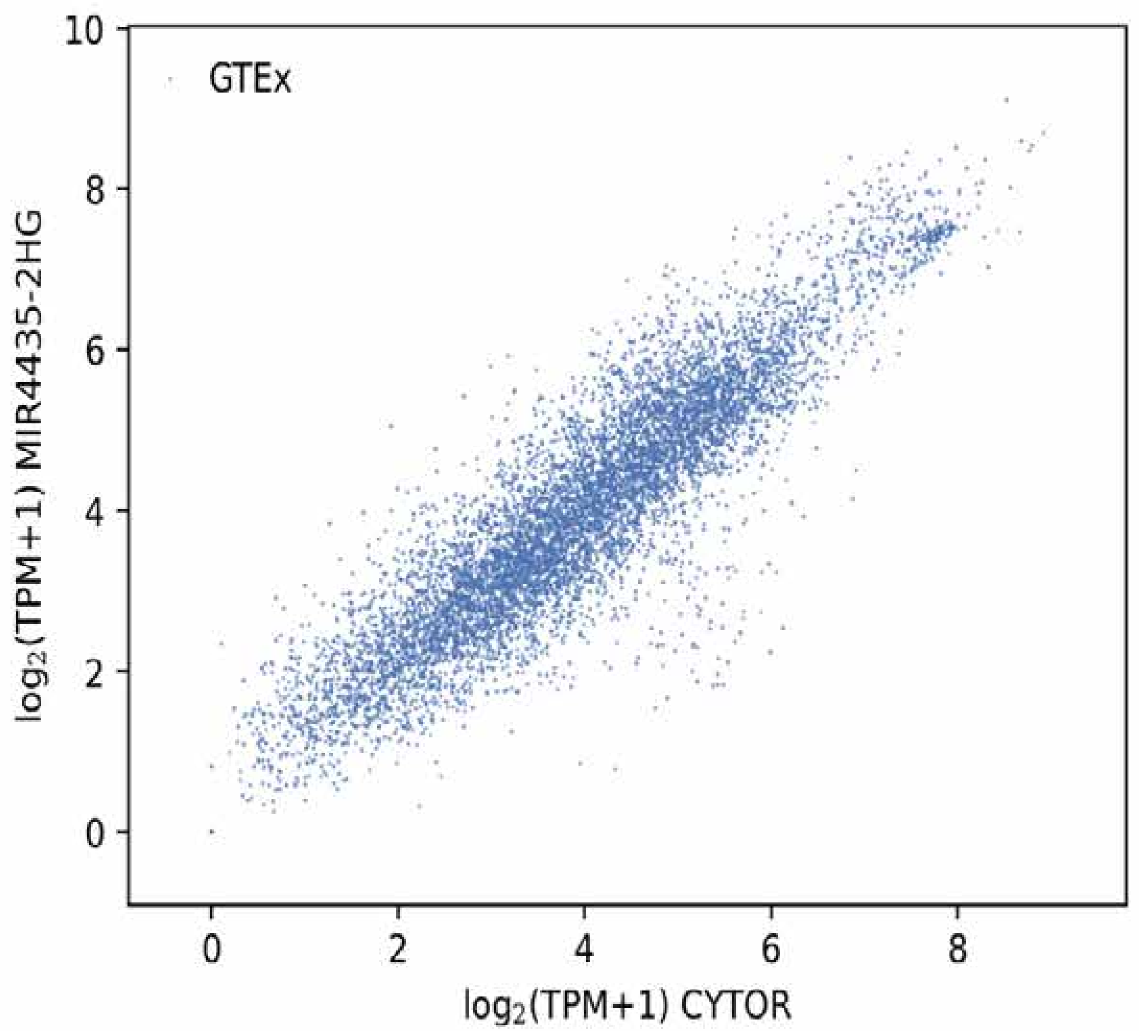


d

MORRID expression profile across tumor samples and paired normal tissues

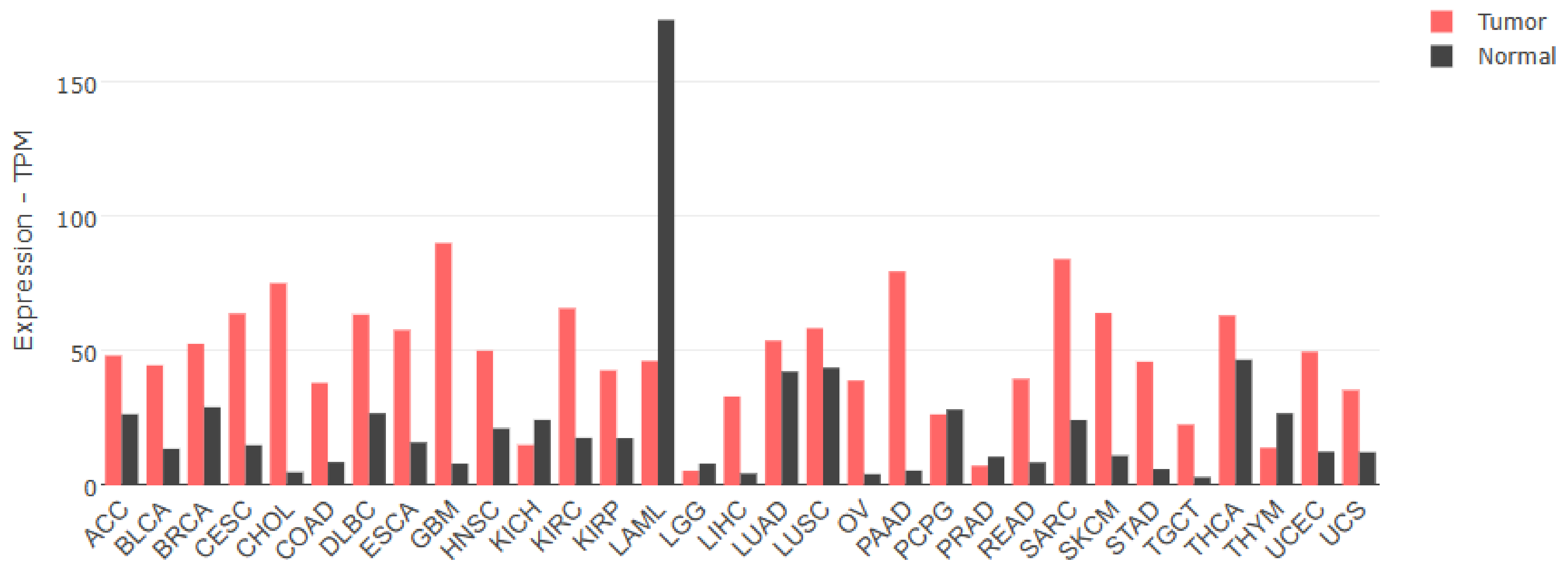


e

CYTOR expression profile across tumor samples and paired normal tissues

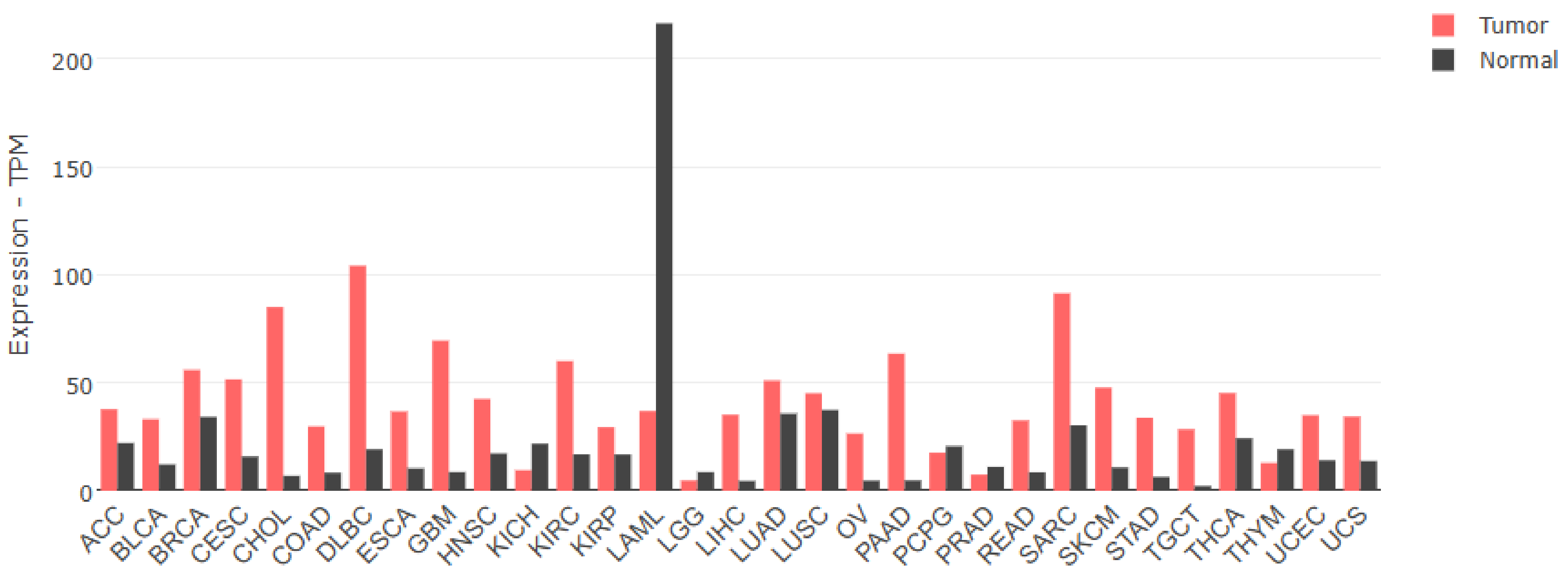


f

Correlation analysis result

| name | name | r | p-value | n_tumor | n_peritumor | n_gtex |
|---|---|---|---|---|---|---|
| CYTOR | MIR4435-2HG | 0.873 | 0 | 9807 | 0 | 0 |

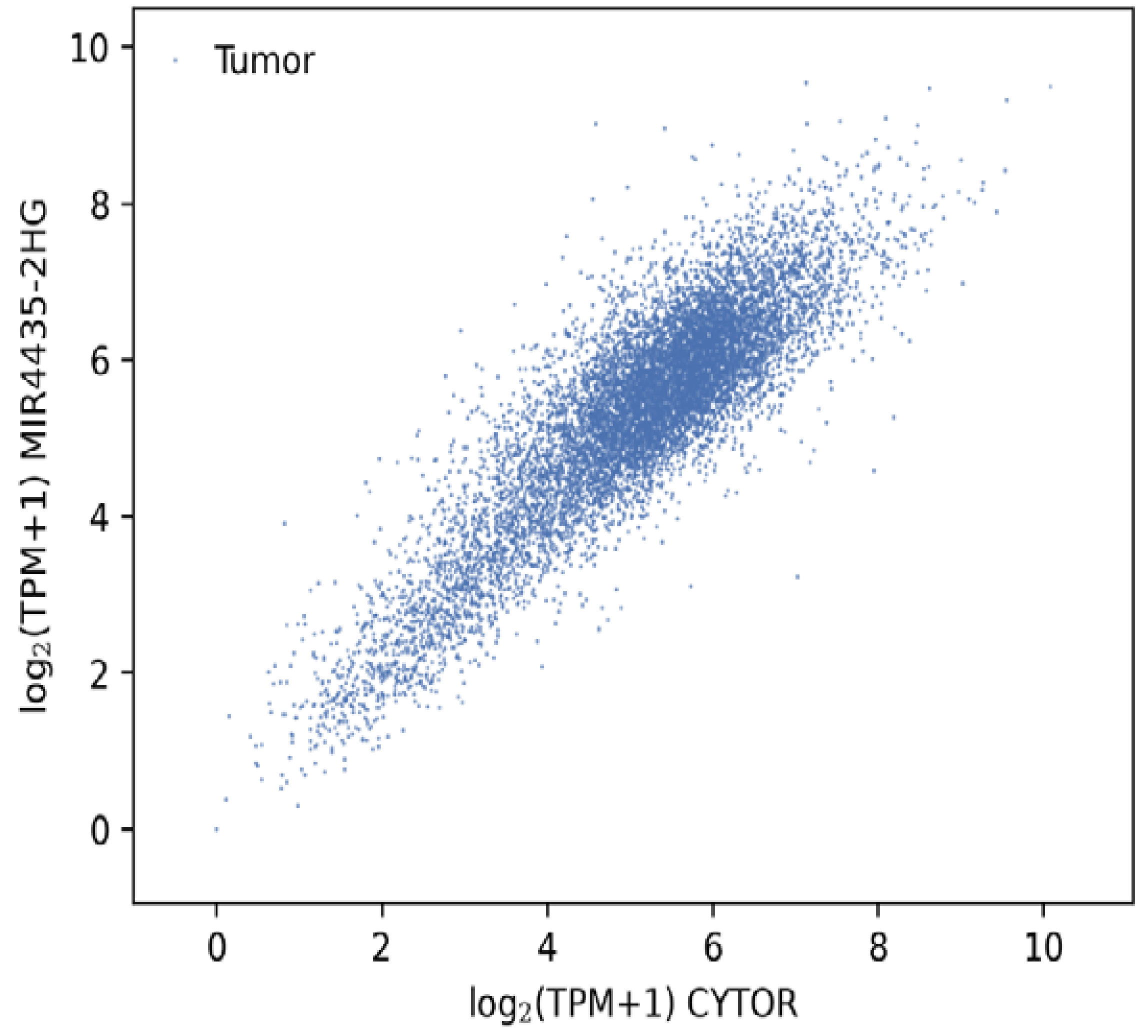
