## Supplementary tables 2 for "Paralogous lncRNAs CYTOR and MORRBID share a conserved trans acting function in MEK ERK signaling"

### **Table S2. Primer sequences**

| Target gene | Direction | Sequence (5′–3′) | Application |
| --- | --- | --- | --- |
| CYTOR/MIR4435-2HG (MORRBID) | Forward | GACATTCCAGACAAGCGGTG | qPCR |
| CYTOR/MIR4435-2HG (MORRBID) | Reverse | TTGCTTGTCAAGGAGAGGCT | qPCR |
| KRCC1 | Forward | TGCATTATGAGATGAGTGAGAGT | qPCR |
| KRCC1 | Reverse | TCAGGCTGAGAATTTGGTGG | qPCR |
| KDM3A | Forward | GAGCTCCACATCAGGTTCAT | qPCR |
| KDM3A | Reverse | TTCAGCCACTTTGATGCAGC | qPCR |
| PLGLB2 | Forward | GACACACTTTCTGGGCACTG | qPCR |
| PLGLB2 | Reverse | GGGCTCTCCTTGACCTGATT | qPCR |
| RGPD1 | Forward | AGTGGTCAAGAAATTGCCTGT | qPCR |
| RGPD1 | Reverse | TCGAGTTCCTGCATGACTGA | qPCR |
| RGPD2 | Forward | CGTCGCCTGGAAAGAAGTTG | qPCR |
| RGPD2 | Reverse | GGATCCCTCTCTTGCACACT | qPCR |
| RMND5A | Forward | CCCCTCTCAGTGTCAGTTTCT | qPCR |
| RMND5A | Reverse | CTGCCTCTGTTCAATCACGG | qPCR |
| BCL2L11 | Forward | GCTTACTATGCAAGGAGGGTATT | qPCR |
| BCL2L11 | Reverse | TGCTGGGTCTTGTTGGTTTG | qPCR |
| BUB1 | Forward | CTGGCTTGGCACTGATTGAC | qPCR |
| BUB1 | Reverse | ACCCCAAAGTAATCGATCTGGT | qPCR |
| LIMS4 | Forward | GTGCCAAGTGTGAGAAACCC | qPCR |
| LIMS4 | Reverse | GAGCAGAGACCACATCACCT | qPCR |
| MERTK | Forward | AGACTGCCTGGATGAACTGTAT | qPCR |
| MERTK | Reverse | TAACGTCTGCTTGGTTCCGA | qPCR |
| RGPD6 | Forward | TGTTACAGGCGTTCAGTGGA | qPCR |
| RGPD6 | Reverse | TTTTGCTGCTCTTTCGACCC | qPCR |
| TMEM87B | Forward | CAGATTGGATGGAACGCTGG | qPCR |
| TMEM87B | Reverse | AGGCATATCTCTGATTGTTTGCT | qPCR |
| FOSL1 | Forward | CAGGCGGAGACTGACAAACTG | qPCR |
| FOSL1 | Reverse | TCCTTCCGGGATTTTGCAGAT | qPCR |
| EPHA4 | Forward | CAGGCGGAGACTGACAAACTG | qPCR |
| EPHA4 | Reverse | TCCTTCCGGGATTTTGCAGAT | qPCR |
| ANTXR1 | Forward | ATTTCAAGTTGTCGTGAGAGGAAAC | qPCR |
| ANTXR1 | Reverse | GACCGAGTCATTGATCTTGAAGCT | qPCR |
| LOXL2 | Forward | GGGTGGAGGTGTACTATGATGG | qPCR |
| LOXL2 | Reverse | CTTGCCGTAGGAGGAGCTG | qPCR |
| CD36 | Forward | TGCAAGTCCTGATGTTTCAGA | qPCR |
| CD36 | Reverse | TGGCTTGACCAATAGGTTGAC | qPCR |
| COL6A3 | Forward | AAGCCTGTGTATCGTGGAG | qPCR |
| COL6A3 | Reverse | TTAAGGCATTGGTCCCAAC | qPCR |
| VANGL2 | Forward | GAGGGCCAGGCTTGTAGTG | qPCR |
| VANGL2 | Reverse | TTTCTGCTCCTCTTCCTGCA | qPCR |
| FAT1 | Forward | GGTCCAGATCGAGGCATTTGA | qPCR |
| FAT1 | Reverse | TCATCTTGCTGTTCTCGGTCTAG | qPCR |
| LIN7A | Forward | ACCCTGGACAGAGGTGTATCA | qPCR |
| LIN7A | Reverse | ACTGTTGCCTTTGCTGTTGC | qPCR |
| MMP13 | Forward | TCACCAATTCCTGGGAAGTCT | qPCR |
| MMP13 | Reverse | TCAGGAAACCAGGTCTGGAG | qPCR |
| NPNT | Forward | TGGGGACAGTGCCAACCTTTCT | qPCR |
| NPNT | Reverse | TGTGCTTACAGGGCCGAGGCT | qPCR |
| SCNN1G | Forward | CGGGAATCAATGCCATTCAGG | qPCR |
| SCNN1G | Reverse | ACACTCCATCAAAGAAGCAGGT | qPCR |
| SPOCD | Forward | CCCCATGGAGTGAAGCTTGT | qPCR |
| SPOCD | Reverse | GCACCATGGGCCTTTTCTTC | qPCR |

### **Table S3. Antibodies**

| Target | Antibody | Catalog number | Company | Application |
| --- | --- | --- | --- | --- |
| MEK2 | MEK2 antibody | Ab265586 | Abcam | Western blot |
| Phospho-MEK1/2 | Phospho-MEK1/2 antibody | #9121 | Cell Signaling Technology | Western blot |
| ERK1/2 | p44/42 MAPK (ERK1/2) antibody | #9102 | Cell Signaling Technology | Western blot |
| Phospho-ERK1/2 | Phospho-p44/42 MAPK (ERK1/2) antibody | #9101 | Cell Signaling Technology | Western blot |
| GAPDH | GAPDH antibody | #2118 | Cell Signaling Technology | Western blot |
| Rabbit IgG | Goat anti-rabbit (HRP-linked) secondary antibody | #7074 | Cell Signaling Technology | Western blot |
| MEK2 | MEK2 antibody (A-1) | sc-13159 | Santa Cruz Biotechnology | CLIP–qPCR |

### **Table S4. Plasmids**

| Plasmid | Source | Identifier | Application |
| --- | --- | --- | --- |
| dCas9-KRAB-mCherry-ZIM3 | Addgene | Plasmid #154473 | Stable CRISPRi cell line generation |
| pSB700-Blasticidin | Addgene | Plasmid #64046 | sgRNA cloning and lentiviral transduction |
| N174 expression vector | Addgene | Plasmid #81061 | EPHA4 overexpression construct generation |

### **Table S5. sgRNAs**

| sgRNA name | Target | Sequence (5′–3′) | Application |
| --- | --- | --- | --- |
| sgRNA1 | CYTOR | GAGGGAAATAAATGACTGGA | CRISPRi knockdown |
| sgRNA3 | CYTOR | GTGTACATCATTGGGAATGG | CRISPRi knockdown |
| NC10010 | Non-targeting control | GTCCACCCTTATCTAGGCTA | Negative control |
